# Ndel1 modulates dynein activation in two distinct ways

**DOI:** 10.1101/2023.01.25.525437

**Authors:** Sharon R Garrott, John P Gillies, Aravintha Siva, Saffron R Little, Rita El Jbeily, Morgan E DeSantis

**Author notes:** indicates corresponding author Correspondence. indicates co-first authors (Sharon R Garrott, John P Gillies).

## Abstract

Dynein is the primary minus-end-directed microtubule motor [1]. To achieve activation, dynein binds to the dynactin complex and an adaptor to form the “activated dynein complex” [2, 3]. The protein Lis1 aids activation by binding to dynein and promoting its association with dynactin and adaptor [4, 5]. Ndel1 and its orthologue Nde1 are dynein and Lis1 binding proteins that help control where dynein localizes within the cell [6]. Cell-based assays suggest that Ndel1/Nde1 also work with Lis1 to promote dynein activation, although the underlying mechanism is unclear [6]. Using purified proteins and quantitative binding assays, we found that Ndel1’s C-terminal region contributes to binding to dynein and negatively regulates binding to Lis1. Using single-molecule imaging and protein biochemistry, we observed that Ndel1 inhibits dynein activation in two distinct ways. First, Ndel1 disfavors the formation of the activated dynein complex. We found that phosphomimetic mutations in Ndel1’s C-terminal domain increase its ability to inhibit dynein-dynactin-adaptor complex formation. Second, we observed that Ndel1 interacts with dynein and Lis1 simultaneously and sequesters Lis1 away from its dynein binding site. In doing this, Ndel1 prevents Lis1-mediated dynein activation. Our work suggests that *in vitro*, Ndel1 is a negative regulator of dynein activation, which contrasts with cellular studies where Ndel1 promotes dynein activity. To reconcile our findings with previous work, we posit that Ndel1 functions to scaffold dynein and Lis1 together while keeping dynein in an inhibited state. We speculate that Ndel1 release can be triggered in cellular settings to allow for timed dynein activation.

## Introduction

Cytoplasmic dynein-1 (dynein) is a microtubule-associated molecular motor that is responsible for nearly all minus-end directed force-generation in most eukaryotes [1]. Dynein traffics hundreds of unique types of cargos, positions the centrosome, facilitates spindle focusing and alignment during mitosis, and strips Spindle Assembly Checkpoint components present at the kinetochore to promote metaphase-anaphase transition [1, 7–10]. Mutations in dynein or its regulatory partners are associated with a host of neurodevelopmental and neurodegenerative diseases [11].

Dynein is a large protein complex comprised of six different subunits, each present in two copies (Figure 1A; Figure S1A). The largest subunit, called the heavy chain, contains a motor domain that is a ring of six AAA (ATPase Associated with various cellular Activities) domains (Figure S1A). AAA1, AAA3, and AAA4 are active ATPase modules [12, 13]. The heavy chain also contains dynein’s microtubule binding domain and a large projection called the tail which acts as a platform for the assembly of the other subunits, including the intermediate chains, the light intermediate chains, and three different light chains (Figure 1A; Figure S1A) [12, 13]. In the absence of other protein factors, dynein exists in an autoinhibited conformation (called “Phi”) and cannot engage productively with the microtubule track (Figure S1A) [14]. Dynein is activated by binding to the multi-subunit protein complex, dynactin, and one of a family of activating adaptor proteins (adaptors) (Figure S1A) [2, 3]. This complex, which we will call the “activated dynein complex”, positions the dynein motor domains in a parallel conformation that is competent to move processively along the microtubule (Figure S1A) [15–18]. In addition to assembling into a complex to activate dynein motility, adaptors also link dynein to cargo [1]. Another regulatory protein, Lis1, promotes formation of the activated dynein complex by binding and converting Phi into a conformation that is primed to bind dynactin and adaptor (Figure S1A) [4, 5, 19–25]. Together, dynactin, adaptors, and Lis1 are three out of four of dynein’s core regulatory machinery.

**Figure 1.**
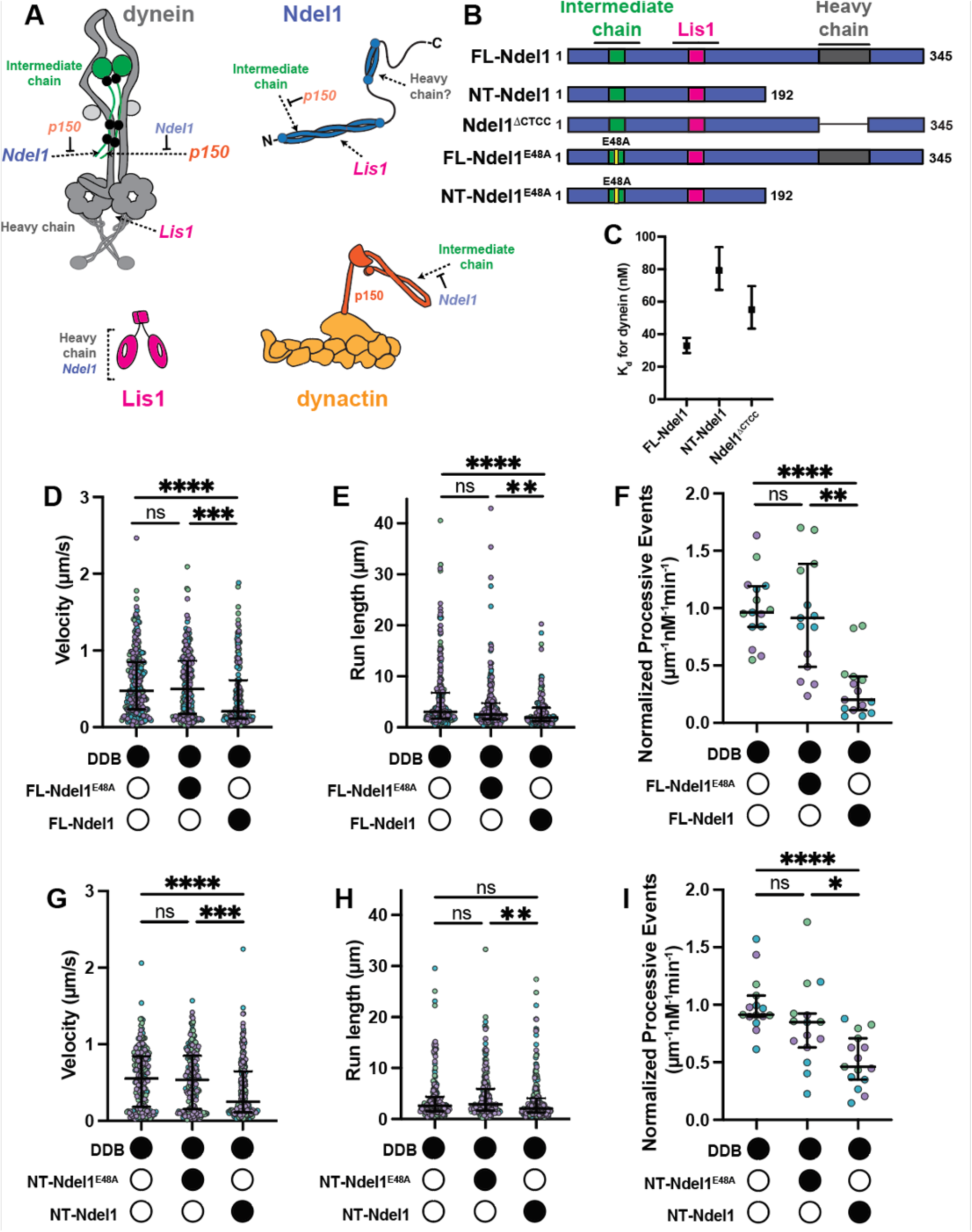
Ndel1 reduces motility of DDB complexes. **A**. Diagrams of dynein and its regulators. Dashed arrows indicate binding sites and blunted arrows indicate competitive binding. Dynein heavy chain (grey) and intermediate chain (green) are labeled. Light intermediate chains (light grey) and light chains (black) are unlabeled **B**. Schematics of Ndel1 constructs. **C**. Kds between dynein and Ndel1 constructs. Error bars indicate 95% confidence intervals. **D**. Single-molecule velocity of DDB complexes in the absence (white circles) or presence (black circles) of 300nM FL-Ndel1^E48A^ or FL-Ndel1. n = 257 (no Ndel1); 209 (FL-Ndel1^E48A^); 181 (FL-Ndel1). Error bars are median±interquartile range. Statistical analysis was performed using a Kruskal-Wallis with Dunn’s multiple comparisons test. p values: ns >0.9999; *** = 0.0001; **** <0.0001. **E**. Single-molecule run lengths of DDB complexes in the absence (white circles) or presence (black circles) of 300nM FL-Ndel1^E48A^ or FL-Ndel1. n = 257 (no Ndel1); 209 (FL-Ndel1^E48A^); 181 (FL-Ndel1). Error bars are median±interquartile range. Statistical analysis was performed using a Kruskal-Wallis with Dunn’s multiple comparisons test. p values: ns =0.1196; ** = 0.0080; **** <0.0001. **F**. Single-molecule events per µm of microtubule per nM dynein for DDB complexes in the absence (white circles) or presence (black circles) of 300nM FL-Ndel1^E48A^ or FL-Ndel1. Data is normalized to the no Ndel1 control for each replicate. n = 15 microtubules per condition. Error bars are median±interquartile range. Statistical analysis was performed using a Brown-Forsythe and Welch ANOVA with Dunnet’s T3 multiple comparisons test. p values: ns >0.9999; ** = 0.0012; **** < 0.0001. **G**. Single-molecule velocity of DDB complexes in the absence (white circles) or presence (black circles) of 300nM NT-Ndel1^E48A^ or NT-Ndel1. n = 193 (no Ndel1); 207 (NT-Ndel1^E48A^); 215 (NT-Ndel1). Error bars are median±interquartile range. Statistical analysis was performed using a Kruskal-Wallis with Dunn’s multiple comparisons test. p values: ns >0.9999; *** = 0.0005; **** <0.0001. **H**. Single-molecule run lengths of DDB complexes in the absence (white circles) or presence (black circles) of 300nM NT-Ndel1^E48A^ or NT-Ndel1. n = 193 (no Ndel1); 207 (NT-Ndel1^E48A^); 215 (NT-Ndel1). Error bars are median±interquartile range. Statistical analysis was performed using a Kruskal-Wallis with Dunn’s multiple comparisons test. p values: ns (No Ndel1 vs NT-Ndel1^E48A^) =0.8699; ns (No Ndel1 vs NT-Ndel1) =0.0960; ** =0.0032. **I**. Single-molecule events per µm of microtubule per nM dynein for DDB complexes in the absence (white circles) or presence (black circles) of 300nM NT-Ndel1^E48A^ or NT-Ndel1. Data is normalized to the no Ndel1 control for each replicate. n = 15 microtubules per condition. Error bars are median±interquartile range. Statistical analysis was performed using a Brown-Forsythe and Welch ANOVA with Dunnet’s T3 multiple comparisons test. p values: ns =0.3462; * = 0.0261; **** < 0.0001.

The fourth member of dynein’s core regulatory partners is the protein Ndel1 and its orthologue, Nde1. Although there is no structure of full-length Ndel1 or Nde1, structures of the N-terminal half of Ndel1 reveal that it is a long coiledcoil [26]. The C-terminal half is largely disordered, except for a short stretch of ∼40 amino acids that forms a coiled-coil (Figure S1B, C) [27]. Many proteins bind to Ndel1/Nde1 to localize dynein throughout the cell cycle. For example, interactions between CENP-F and Ndel1/Nde1 help position dynein at the nuclear pore for centrosome positioning and at the kinetochore, where dynein will eventually traffic checkpoint proteins toward the spindle poles to facilitate the metaphase-anaphase transition [10, 28–30]. Ndel1 and Nde1 may also support dynein localization to cargo in interphase. For example, interactions between Ndel1/Nde1 and Rab9 promotes dynein localization to endosomes [31].

In addition to localizing dynein within the cell, there is evidence that Ndel1 and Nde1 directly support dynein activation. In human cells, concurrent depletion of Ndel1 and Nde1 cause Golgi dispersal, while inhibition of Ndel1 and Nde1 by a function-blocking antibody cause acidic cargo dispersal [8, 32]. Both effects suggest a reduction in dynein-driven minus-end directed trafficking. Depletion of the Ndel1 homologue in fungi causes defects in nuclear positioning in a manner that is consistent with reduced dynein activity [33–36]. Furthermore, in filamentous fungi, expression of a dynein mutant that cannot assume the Phi structure can partially rescue the nuclear positioning defect caused by Ndel1 depletion, suggesting that Ndel1 serves to activate dynein [24].

Ndel1 and Nde1 likely work together with Lis1 to support dynein activity. *In vitro* experiments conducted with purified mammalian and yeast proteins show that Ndel1/Nde1 promote Lis1-dynein association [37, 38]. Additionally, cell-based assays performed in many organisms and cell types show that deleterious effects of knockdown of either protein can be rescued by overexpression of the other, suggesting that Lis1 and Ndel1/Nde1 operate in the same pathway. For example, Ndel1 depletion in filamentous fungi and *Xenopus* extracts results in nuclear distribution defects and spindle focusing defects, respectively. Both phenotypes are rescued by Lis1 overexpression [39–41]. Similarly, in developing fly brains, dendritic arborization defects caused by Nde1 deletion are rescued by Lis1 overexpression [42]. Conversely, in mammalian cells, Ndel1 expression rescues spindle orientation defects and Golgi dispersal caused by Lis1 knockdown [8, 43]. A prevailing model is that Ndel1/Nde1 tether dynein and Lis1, stabilizing their interaction [37, 39, 44]. A prediction from this model is that Ndel1/Nde1 will increase Lis1’s ability to form the activated dynein complex.

Multiple structural and biochemical studies have refined our understanding of how dynactin, adaptors, and Lis1 interact with dynein to facilitate activation (Figure S1A) [4, 5, 15–21,23, 25]. Ndel1 and Nde1’s position in the interaction network of dynein and its regulators is less clear. In addition to binding Lis1’s beta propeller, Ndel1 and Nde1 bind to the dynein intermediate chain and heavy chain (Figure 1A). The first ∼50 amino acids of Ndel1/Nde1 interacts with the disordered N-terminal tail of the intermediate chain (Figure 1A) [38, 39, 44–47]. This interaction has been validated *in vitro* and in cell-based assays. Interestingly, Ndel1/Nde1 bind the intermediate chain competitively with a coiled-coil in dynactin’s p150 subunit (called CC1) (Figure 1A) [46–49]. Ndel1’s purported heavy chain-binding site, which is less validated, is within the C-terminal coiled-coil of Ndel1 (amino acids ∼253-293) (Figure 1A) [50, 51]. On the dynein side, this interaction has not been mapped with high confidence but may span AAA1 and the heavy chain C-terminal tail [51].

We set out to investigate how Ndel1 affected dynein activity and test the model that Ndel1 tethers Lis1 to dynein to promote activation. Using single-molecule imaging and protein biochemistry, we found that Ndel1 disfavors formation of the activated dynein complex, most likely via competition with p150 for intermediate chain-binding. Despite Ndel1’s C-terminus being dispensable for inhibiting dynein activation, it increases the potency of inhibition. Further, phosphomimetic mutations in Ndel1’s C-terminus increase its ability to inhibit complex formation. Our work also revealed that Ndel1 and Lis1 binding is negatively regulated by the C-terminal half of Ndel1, suggesting that Ndel1 is autoinhibited with respect to Lis1 association. We found that although Ndel1 can bind both Lis1 and dynein simultaneously, Lis1 cannot bind Ndel1 and dynein at the same time. This competition significantly attenuates Lis1’s ability to promote activated dynein complex assembly in the presence of Ndel1. Our work suggests that Ndel1 has two distinct modes for preventing dynein activation: disfavoring dynein complex assembly by binding the intermediate chain and preventing Lis1-mediated activation.

## Results

### The N-terminal half of Ndel1 inhibits dynein motility

The function of Ndel1’s two dynein binding sites has been controversial. To characterize the interaction between dynein and Ndel1, we purified SNAP-tagged, recombinant full-length human dynein with all associated accessory chains (dynein), full-length Ndel1 (FL-Ndel1), and several Ndel1 mutant or deletion constructs (Figure 1B; Figure S1D). These constructs include a truncation of Ndel1 containing only the N-terminal coiled-coil (NT-Ndel1), a well-characterized Ndel1 mutant that disrupts binding to the intermediate chain in both the FL-Ndel1 and NT-Ndel1 backgrounds (FL-Ndel1^E48A^ and NT-Ndel1^E48A^), and a construct with the coiled-coil containing the purported heavy chain-binding site deleted from Ndel1’s C-terminal region (Ndel1^.6.CTCC^) (Figure 1B)[39, 44, 45, 51]. All Ndel1 constructs had HaloTags at their N-termini and 6X-His tags at their C-termini [52].

First, we determined the binding affinities between each Ndel1 construct and dynein to determine the relative contribution of each of Ndel1’s dynein binding sites. To do this, we conjugated increasing amounts of Ndel1 to magnetic beads via the HaloTag and quantified dynein depletion via SDS-PAGE. We found that FL-Ndel1 bound dynein with a K_d_ of ∼33 nM (Figure 1C; Figure S1E). Both NT-Ndel1 and Ndel1.6.CTCC displayed a reduced affinity for dynein (K_d_ of 79 nM and 55 nM, respectively) (Figure 1C; Figure S1F, G). Together, these results suggest that regions in Ndel1’s N-terminal and C-terminal halves both contribute to interaction with dynein. FL-Ndel1^E48A^ did not appreciably bind dynein (Figure S1H), which indicates that the interaction between Ndel1’s C-terminal half and dynein is not sufficiently strong to promote binding in the absence of a productive interaction between Ndel1 and the intermediate chain.

Previous work suggested that mammalian Ndel1 reduces dynein’s affinity for microtubules [38, 47, 53, 54]. Using single-molecule total internal reflection fluorescence microscopy (smTIRF), we monitored the microtubule binding activity of TMR-labeled dynein in the absence and presence of FL-Ndel1. We observed that FL-Ndel1 did not alter the density of single molecules of dynein on microtubules, indicating that Ndel1 does not modulate dynein’s microtubule binding affinity (Figure S1I). This finding is consistent with what has been observed with yeast proteins [37].

Next, we investigated if Ndel1 affects the motility of activated dynein complexes. To do this, we assembled TMR-or Alexa647-labeled dynein, purified dynactin, and the purified, recombinant adaptor, BicD2 into complexes (DDB) (Figure S1D). Because many full-length adaptors display autoinhibition, we used a well-characterized truncation of BicD2 containing amino acids 25-398 (hereafter called BicD2) [4, 55–57]. We next incubated DDB with FL-Ndel1 or FL-Ndel1^E48A^ for ten minutes to allow samples to reach equilibrium, then imaged motility of DDB using smTIRF. We observed that FL-Ndel1, but not FL-Ndel1^E48A^, significantly reduced the velocity, run length, and landing rate of processive, motile events compared to DDB alone (Figure 1D-F; Figure S2A). We did not observe FL-Ndel1 co-migrating with processive events, suggesting that Ndel1 does not alter motility by remaining bound to the moving complex (Figure S2B).

To determine the relative contribution of Ndel1’s C-terminal binding site to its effect on dynein motility, we repeated the smTIRF motility experiments with NT-Ndel1 and NT-Ndel1^E48A^ (Figure S2C). As expected, NT-Ndel1^E48A^ did not affect any motility parameters (Figure 1G-I). Interestingly, NT-Ndel1 had the same effect on velocity as FL-Ndel1, showing a ∼2-fold reduction compared to DDB alone (Figure1D, G). NT-Ndel1 also reduced the landing rate of processive events, but to a lesser extent than FL-Ndel1 (compare a ∼5-fold reduction in landing rate with FL-Ndel1 to a ∼2-fold reduction with NT-Ndel1) (Figure 1F, I). NT-Ndel1 did not significantly reduce the run length of motile events compared to DDB alone (Figure 1H).

Together, these data suggest that Ndel1 reduces the number of activated dynein complexes as well as the velocity and run length of processive DDB. Because both NT-Ndel1 and FL-Ndel1 reduced velocity and total processive events, we reason that the inhibitory effect of Ndel1 is mediated by its N-terminal intermediate chain-binding site. The increased inhibition observed with FL-Ndel1 compared to NT-Ndel1 is consistent with the K_d_ measurements that show that Ndel1’s C-terminal half increases its association with dynein (Figure 1C). Thus, our data suggest that the C-terminal region of Ndel1 increases the potency of inhibition, but that Ndel1’s N-terminal coiled-coil is necessary and sufficient.

### Ndel1 competes with dynactin and adaptor for dynein binding

We next asked if Ndel1 had an inhibitory effect on dynein activated with different adaptors. Here, we reconstituted dynein motility with the adaptor ninein-like (NINL), and dynactin (DDN) and determined if FL-Ndel1 affected DDN motility (Figure 2A-C; Figure S2D; Figure S1D). As with BicD2, we used a truncation of NINL (aa 1-702) to avoid autoinhibition [4, 58, 59]. As was observed with DDB, FL-Ndel1 reduced the landing rate of processive DDN events (Figure 2C). Unlike with DDB, however, FL-Ndel1 had little effect on the velocity or run length of processive DDN complexes (Figure 2A, B).

**Figure 2.**
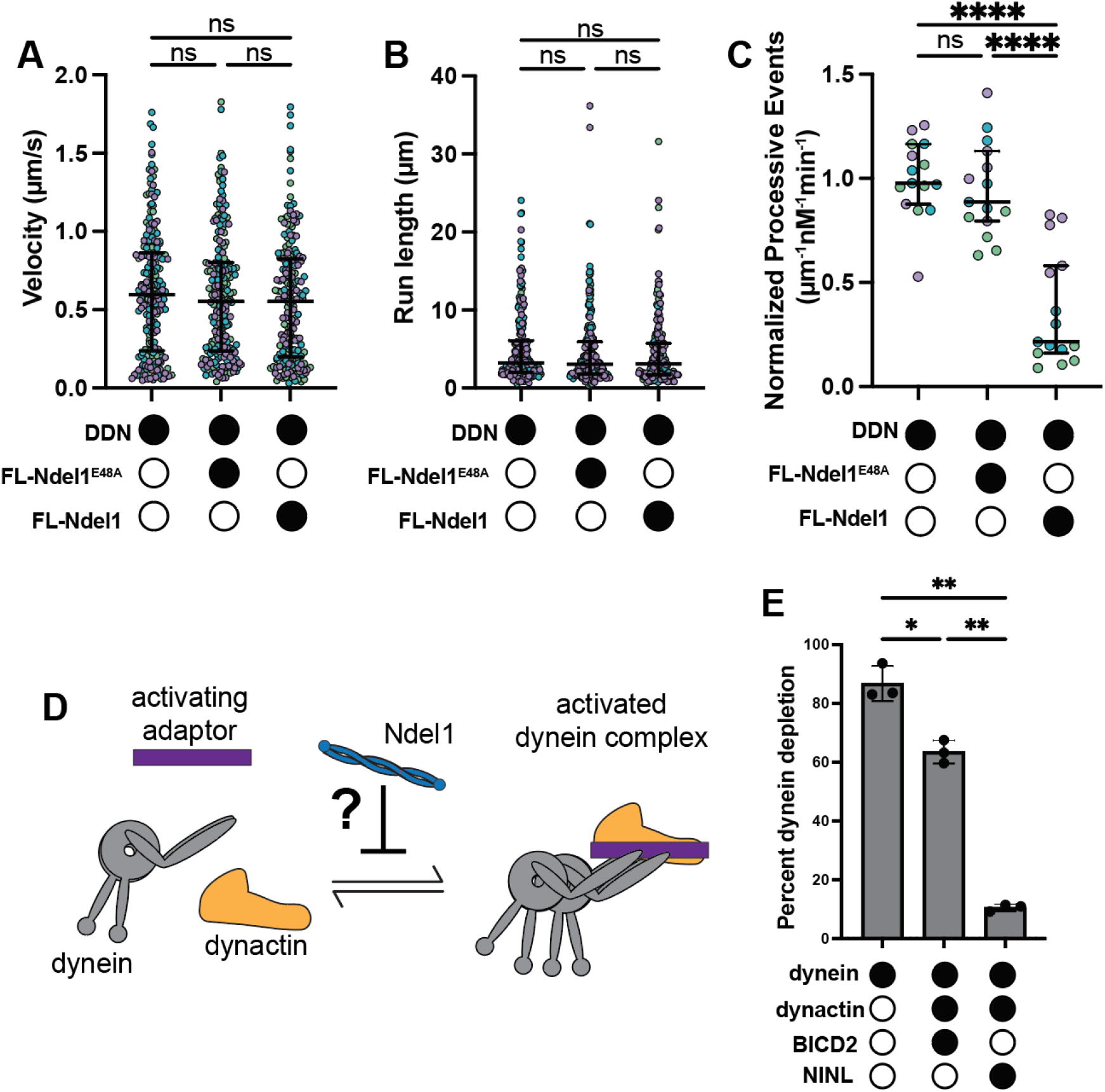
Ndel1 differentially effects activated complexes formed with different adaptors. **A**. Single-molecule velocity of DDN complexes in the absence (white circles) or presence (black circles) of 300nM FL-Ndel1^E48A^ or FL-Ndel1. n = 214 (no Ndel1); 203 (FL-Ndel1^E48A^); 194 (FL-Ndel1). Error bars are median±interquartile range. Statistical analysis was performed using a Kruskal-Wallis with Dunn’s multiple comparisons test. p values: no Ndel1 vs FL-Ndel1 = 0.4462; others are >0.9999. **B**. Single-molecule run lengths of DDN complexes in the absence (white circles) or presence (black circles) of 300nM FL-Ndel1^E48A^ or FL-Ndel1. n = 214 (no Ndel1); 203 (FL-Ndel1^E48A^); 194 (FL-Ndel1). Error bars are median±interquartile range. Statistical analysis was performed using a Kruskal-Wallis with Dunn’s multiple comparisons test. p values: no Ndel1 vs FL-Ndel1 = 0.2971; no Ndel1 vs FL-Ndel1^E48A^ = 0.9669; FL-Ndel1^E48A^ vs FL-Ndel1 > 0.9999. **C**. Single-molecule events per µm of microtubule per nM dynein for DDN complexes in the absence (white circles) or presence (black circles) of 300nM FL-Ndel1^E48A^ or FL-Ndel1. Data is normalized to the no Ndel1 control for each replicate. n = 15 microtubules per condition. Error bars are median±interquartile range. Statistical analysis was performed using a Brown-Forsythe and Welch ANOVA with Dunnet’s T3 multiple comparisons test. p values: ns = 0.8528; **** <0.0001. **D**. Schematic of the assay to test if Ndel1 affects formation of activated dynein complexes. **E**. Percentage of dynein bound to FL-Ndel1 conjugated beads in the absence (white circles) or presence (black circles) of dynactin, BicD2 and NINL. Statistical analysis was performed using a Brown-Forsythe and Welch ANOVA with Dunnet’s T3 multiple comparisons test. p values: * = 0.0259; ** (dynein vs DDN) = 0.0044; ** (DDB vs DDN) = 0.0042.

We reasoned that the reduction in processive events caused by Ndel1 with both adaptors could be explained if Ndel1 acted upstream of activation and disfavored the formation of the activated dynein complex (Figure 2D). To test this hypothesis, we conjugated FL-Ndel1 via the C-terminal 6X-His-tag to magnetic beads and monitored its ability to bind and deplete dynein from solution. Here, we incubated FL-Ndel1-beads with dynein alone; dynein, dynactin, and BicD2; or dynein, dynactin, and NINL. FL-Ndel1 depleted over 80% of the dynein in solution in the absence of dynactin and adaptor (Figure 2E). Inclusion of dynactin and BicD2 resulted in less binding between dynein and Ndel1, with a depletion of 65% of the dynein (Figure 2E). Remarkably, only ∼11% of the total dynein was depleted by FL-Ndel1 in the presence of dynactin and NINL (Figure 2E). Because dynactin and either adaptor reduced dynein’s binding affinity for FL-Ndel1, these results suggest that Ndel1 competes with dynactin and adaptors for dynein binding. This result supports the model that Ndel1 disfavors complex formation. These results also highlight that DDN is more refractory to Ndel1 inhibition than DDB, which may explain why DDN velocity and run length are less affected by Ndel1 than DDB.

### Phosphomimetic Ndel1 mutations enhance dynein inhibition by Ndel1

Ndel1 and its orthologue Nde1 are phosphoproteins [60]. Cell-based studies have revealed that phosphorylation of Ndel1 and Nde1 by many different kinases modulates their sub-cellular localization or their ability to bind dynein or Lis1 [61–65]. Many kinases target regions in Ndel1 and Nde1 in or around the C-terminal coiled-coil that contains the purported heavy chain-binding site (Figure 3A; Figure S3A) [61, 65]. Phosphorylation of Ndel1’s C-terminal region regulates Ndel1’s ability to promote the correct subcellular localization of dynein and/or Lis1. For example, phosphorylation by CDK5 and CDK1 enable the Ndel1-mediated recruitment of dynein or Lis1 to the nuclear pore and kinetochore [61, 62]. Additionally, in numerous cell-based assays, CDK5 phosphorylation of Ndel1 has been shown to positively regulate association with dynein and dynein-driven cargo trafficking, however in vitro experiments do not support this observation [44, 61, 65, 66].

**Figure 3.**
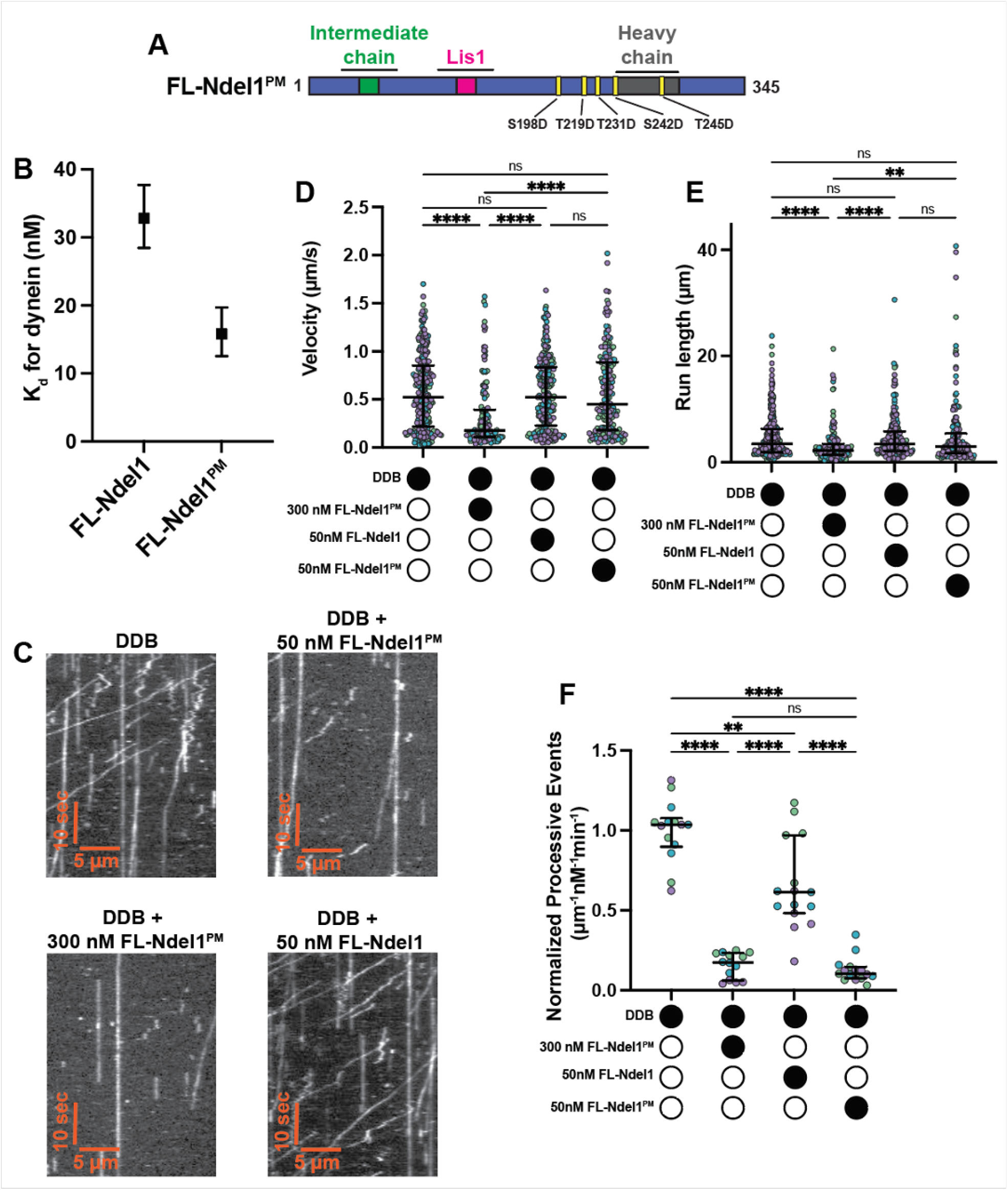
Phosphomimetic mutations in Ndel1’s C-terminal region increase the efficacy of activated dynein complex inhibition. **A**. Schematic of FL-Ndel1^PM^. **B**. K_d_s between dynein and FL-Ndel1 constructs. Error bars indicate 95% confidence intervals. FL-Ndel1 data previously shown in Figure 1C. **C**. Example kymographs of DDB in the presences and absence of FL-Ndel1^PM^ and FL-Ndel1. **D**. Single-molecule velocity of DDB complexes in the absence (white circles) or presence (black circles) of FL-Ndel1^PM^ or FL-Ndel1 at the concentrations indicated. n = 276 (no Ndel1); 178 (50nM FL-Ndel1); 221 (50nM FL-Ndel1^PM^); 137 (300nM FL-Ndel1^PM^). Error bars are median±interquartile range. Statistical analysis was performed using a Kruskal-Wallis with Dunn’s multiple comparisons test. p values: ns > 0.9999; **** <0.0001. **E**. Single-molecule run lengths of DDB complexes in the absence (white circles) or presence (black circles) of FL-Ndel1^PM^ or FL-Ndel1 at the concentrations indicated. n = 276 (no Ndel1); 178 (50nM FL-Ndel1); 221 (50nM FL-Ndel1^PM^); 137 (300nM FL-Ndel1^PM^). Error bars are median±interquartile range. Statistical analysis was performed using a Kruskal-Wallis with Dunn’s multiple comparisons test. p values: ns (no Ndel1 vs 50nM FL-Ndel1^PM^) >0.9999; ns (no Ndel1 vs 50nM FL-Ndel1^PM^) = 0.3266; ns (50nM Ndel1 vs 50nM FL-Ndel1^PM^) = 0.5724; ** = 0.0057; **** <0.0001. **F**. Single-molecule events per µm of microtubule per nM dynein for DDB complexes in the absence (white circles) or presence (black circles) of FL-Ndel1^PM^ or FL-Ndel1 at the concentrations indicated. Data is normalized to the no Ndel1 control for each replicate. n = 15 microtubules per condition. Error bars are median±interquartile range. Statistical analysis was performed using a Brown-Forsythe and Welch ANOVA with Dunnet’s T3 multiple comparisons test. p values: ns = 0.8912; ** = 0.0042; **** <0.0001.

To explore the effect of phosphorylation on Ndel1’s ability to modulate dynein motility, we purified a FL-Ndel1 construct with the five sites phosphorylated by CDK5 (amino acids S198, T219, T231, S242, and T245) mutated to aspartic acid to mimic phosphorylation (FL-Ndel1^PM^) (Figure 3A; Figure S1D). First, we determined that FL-Ndel1^PM^ binds dynein with a 2-fold higher affinity than FL-Ndel1 (Figure 3B; Figure S3B) using the quantitative binding assay outlined above. Next, we determined if FL-Ndel1^PM^ functioned like FL-Ndel1 in DDB motility assays (Figure 3C-F). At the concentration of Ndel1 used in previous smTIRF assays (300 nM), FL-Ndel1^PM^ caused a decrease in the run length of processive events that was comparable to what we observed with FL-Ndel1 (Figure 3C,E; Figure 1E; Figure S2A). FL-Ndel1^PM^ reduced the velocity and landing rate to a slightly greater extent than FL-Ndel1 (Figure 3C, D, F; Figure 1D, F; Figure S2A). To probe how much more effective the FL-Ndel1^PM^ construct was, we reduced the concentration of FL-Ndel1^PM^ and FL-Ndel1 to 50 nM and repeated the smTIRF motility experiments with DDB. At the reduced concentration, neither FL-Ndel1^PM^ nor FL-Ndel1 had a statistically significant reduction in velocity or run-length (Figure 3C-E; Figure 1D, E; Figure S2A). Remarkably, inclusion of 50 nM FL-Ndel1^PM^ reduced the processive landing rate of DDB by 10-fold, while 50 nM FL-Ndel1 reduced the landing rate by only 1.7-fold (Figure 3C, F). We conclude that the phosphomimetic mutations in Ndel1 promote its binding to dynein and thereby increase its ability to inhibit the formation of the activated dynein complex.

### Lis1 interactions with Ndel1 and dynein are mutually exclusive

Ndel1’s ability to bind Lis1 is well-documented, however the structural basis for their interaction is poorly defined [26, 38, 40, 45, 67, 68]. To dissect the molecular determinants of the interaction between Lis1 and Ndel1, we determined the binding affinity between Lis1 and FL-Ndel1, NT-Ndel1, and FL-Ndel1^PM^. We conjugated each construct of Ndel1 to magnetic resin via the N-terminal HaloTag and monitored depletion of Lis1 from solution. We performed the initial binding experiments at 36 mM KCl, which matches the experimental conditions for all previous binding experiments and the smTIRF motility assays. At this ionic strength, the K_d_ between Lis1 and Ndel1 was far too low to accurately measure in the bead-based assay (data not shown). Therefore, we increased the ionic strength of the final assay buffer to 150 mM KCl and repeated the binding experiments. We found that all constructs of Ndel1 bound Lis1 with high affinity. Both FL-Ndel1 and FL-Ndel1^PM^ had a K_d_ of ∼4 nM for Lis1 (Figure 4A; Figure S4A, B). These data suggest that phosphomimetic mutations of Ndel1 at the CDK5 target sites do not affect binding to Lis1 (Figure 3B). Interestingly, NT-Ndel1 exhibited an increased affinity for Lis1, with a K_d_ of ∼1.2 nM (Figure 4A; Figure S4C), which shows that removing Ndel1’s C-terminus increases its association with Lis1. This result is consistent with crosslinking-mass spectrometry data that suggest that Ndel1’s C-terminus can fold back towards the N-terminal coiled-coil and partially occlude the Lis1 binding site [69]. To further validate this model, we used CoLab Fold to predict the structure of full-length Ndel1 [70, 71]. We found that the C-terminal half of Ndel1 (that contains the predicted heavy chain binding site) folds back and interacts with the mapped Lis1 binding site (Figure 4B, Figure S4D). CoLab Fold predictions of just the C-terminal half of Ndel1 (in the absence of the N-terminal portion) indicate that the putative heavy chain binding site assumes a coiled-coil fold (Figure S1B, C). Given the two conformations of Ndel1’s C-terminal region predicted by CoLab Fold, it’s possible that Ndel1 exhibits a conformational equilibrium that regulates binding to Lis1 (Figure S4E).

**Figure 4.**
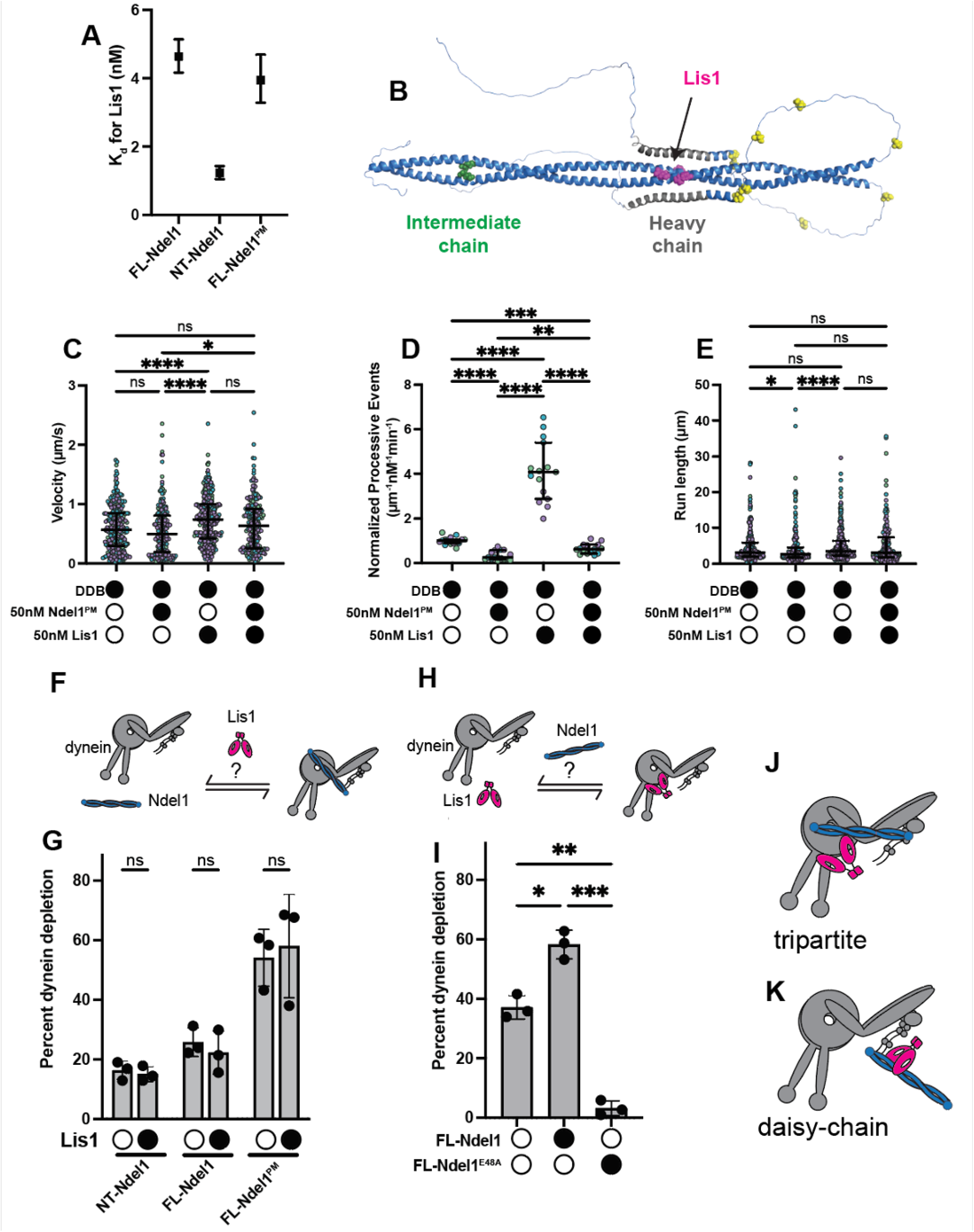
Ndel1 tethers Lis1 and dynein while preventing Lis1-mediated dynein activation. **A**. Kds between Lis1 and Ndel1 constructs. Error bars indicate 95% confidence intervals. **B**. Model generated in ColabFold with the FL-Ndel1 C terminus folded back. Green indicates IC binding region, pink indicates Lis1 binding region, grey indicates heavy chain binding region, and yellow indicates CDK5 phosphorylation sites. **C**. Single-molecule velocity of DDB complexes in the absence (white circles) or presence (black circles) of 50nM FL-Ndel1^PM^ and Lis1. n = 256 (no Ndel1/Lis1); 200 (FL-Ndel1^PM^); 351 (Lis1); 200 (FL-Ndel1^PM^+Lis1). Error bars are median±interquartile range. Statistical analysis was performed using a Kruskal-Wallis with Dunn’s multiple comparisons test. p values: ns (no Ndel1/Lis1 vs. FL-Ndel1^PM^) = 0.3759; ns (no Ndel1/Lis1 vs. FL-Ndel1^PM^+Lis1) > 0.9999; ns (Lis1 vs FL-Ndel1^PM^+Lis1) = 0.0540; * = 0.0172; **** <0.0001. **D**. Single-molecule events per µm of microtubule per nM dynein for DDB complexes in the absence (white circles) or presence (black circles) of 50nM FL-Ndel1^PM^ and Lis1. Data is normalized to the no Ndel1 control for each replicate. n = 15 microtubules per condition. Error bars are median±interquartile range. Statistical analysis was performed using a Brown-Forsythe and Welch ANOVA with Dunnet’s T3 multiple comparisons test. p values: ** = 0.0027; *** = 0.0004; **** <0.0001. **E**. Single-molecule run lengths of DDB complexes in the absence (white circles) or presence (black circles) of 50nM FL-Ndel1^PM^ and Lis1. n = 256 (no Ndel1/Lis1); 200 (FL-Ndel1^PM^); 351 (Lis1); 200 (FL-Ndel1^PM^+Lis1). Error bars are median±interquartile range. Statistical analysis was performed using a Kruskal-Wallis with Dunn’s multiple comparisons test. p values: ns (no Ndel1/Lis1 vs. Lis1) = 0.7162; ns (no Ndel1/Lis1 vs. FL-Ndel1^PM^+Lis1) > 0.9999; ns (FL-Ndel1^PM^ vs FL-Ndel1^PM^+Lis1) = 0.1069; ns (Lis1 vs FL-Ndel1^PM^+Lis1) = 0.5455; * = 0.0367; **** <0.0001. **F**. Schematic of the assay to test if Lis1 affects Ndel1’s ability to bind dynein. **G**. Percent of dynein bound to Ndel1 conjugated beads in the absence (white circles) or presence (black circles) of Lis1. Statistical analysis was performed using a Brown-Forsythe and Welch ANOVA with Dunnet’s T3 multiple comparisons test. p values: NT-Ndel1 = 0.9300; FL-Ndel1 = 0.8589; FL-Ndel1^PM^ = 0.9770. **H**. Schematic of the assay to test if Ndel1 affects Lis1’s ability to bind dynein. **I**. Percent of dynein bound to Lis1 conjugated beads in the absence (white circles) or presence (black circles) of FL-Ndel1 or FL-Ndel1^E48A^. Statistical analysis was performed using a Brown-Forsythe and Welch ANOVA with Dunnet’s T3 multiple comparisons test. p values: * = 0.0106; ** = 0.0026; *** = 0.0009. **J**. Schematics of the tripartite dynein-Lis1-Ndel1 complex. **K**. Schematics of the daisy-chain dynein-Lis1-Ndel1 complex.

*S. cerevisiae* Ndel1 (Ndl1 in yeast) and Lis1 (Pac1 in yeast) have a synergistic effect on Pac1-mediated alteration of dynein velocity [37]. Because yeast dynein exhibits processive motility on its own, previous experiments that tested Ndl1 and Pac1’s effect on dynein were conducted in the absence of dynactin and an adaptor [37, 72]. Inclusion of Pac1 in this context reduces the velocity of dynein, and Ndl1 amplifies this effect [4, 37, 38]. To test if mammalian Lis1 and Ndel1 showed synergistic activity, we performed the smTIRF motility assay with DDB, 50 nM FL-Ndel1^PM^, and/or 50nM Lis1 (Figure 4C-E; Figure S4F). As previously observed, Lis1 alone resulted in DDB complexes that moved with a faster velocity and exhibited a higher landing rate (Figure 4C, D) [4, 5, 22, 23]. At this concentration, Lis1 did not significantly increase the run length of processive events (Figure 4E). Next, we performed the DDB smTIRF motility experiments with equimolar FL-Ndel1^PM^ and Lis1. If Ndel1 tethers Lis1 to dynein as has been proposed, we would anticipate that Lis1 and FL-Ndel1^PM^ should activate dynein motility (i.e. increase velocity, landing rate, and/or run length) to a greater extent than what was observed with Lis1 alone. Instead, we observed that the negative FL-Ndel1^PM^ effect was attenuated by the positive Lis1 effect, with resulting events moving at velocities that were not significantly different than DDB alone (Figure 4C). Remarkably, we observed that the inhibitory effect of FL-Ndel1^PM^ outweighed the activating effect of Lis1 with respect to landing rate. Compared to DDB alone, Lis1 resulted in a ∼4-fold increase in landing rate, while FL-Ndel1^PM^ and Lis1 together exhibited ∼2-fold reduction in landing rate (Figure 4D). These results suggest that Ndel1 and Lis1 do not interact synergistically to increase dynein activation and that the Ndel1-mediated inhibition of DDB assembly prevents Lis1 from promoting complex formation.

We have observed that Lis1 binds Ndel1 directly, and that Ndel1’s N-and C-termini both contribute to dynein binding (Figure 1C). It is also well-established that Lis1 binds dynein’s motor domain at the interface of AAA3 and AAA4 (AAA3/4) and along the stalk that emanates from AAA4 and connects dynein’s ATPase ring to the microtubule-binding domain (Figure S1A) [4, 19, 21]. We set out to dissect if Ndel1, Lis1, and dynein mutually influence each other’s binding. First, we asked if Lis1 affects Ndel1-dynein interaction (Figure 4F, G). We conjugated NT-Ndel1, FL-Ndel1, or FL-Ndel1^PM^ to magnetic beads and monitored the amount of dynein depleted by each Ndel1 construct in the absence and presence of Lis1. Each Ndel1 construct depleted different amounts of dynein from solution, which is consistent with the measured K_d_ values (Figure 1C; Figure 3B). However, we observed that Lis1 had no effect on the ability of any of the Ndel1 constructs to deplete dynein (Figure 4G). This suggests that Lis1 does not regulate the Ndel1-dynein interaction.

Next, we asked if Ndel1 influences the Lis1-dynein interaction (Figure 4H, I). We conjugated Lis1 to magnetic beads via an N-terminal HaloTag. We incubated dynein alone or with tagless FL-Ndel1 (i.e. no HaloTag) and asked if Ndel1 affects the amount of dynein depleted by Lis1. In these experiments, in the absence of FL-Ndel1, we observed ∼40% of the dynein was depleted by Lis1, which is consistent with the Lis1-dynein K_d_ measured (Figure 4I; Figure S4G). Inclusion of FL-Ndel1 increased the amount of dynein depleted by Lis1 to ∼60% (Figure 4I). This suggests that Ndel1 increases the association of dynein and Lis1, which is consistent with previous reports [37, 38].

There are two possible explanations for the apparent increased affinity between Lis1 and dynein in the presence of FL-Ndel1. First, Ndel1, dynein, and Lis1 could assemble in a tripartite structure, with Lis1 bound to dynein at the AAA3/4 and stalk sites while Ndel1 bridges the dynein motor to interact with the intermediate chain and Lis1 simultaneously (Figure 4J). Second, it is possible that Lis1 cannot bind dynein and Ndel1 at the same time and, instead, Ndel1 “daisy-chains” Lis1 to dynein (Figure 4K). This would lead to an increased apparent affinity between Lis1-dynein because Ndel1 binds dynein with a higher affinity than Lis1 does (compare a K_d_ of 33 nM for dynein-Ndel1 to 101 nM for dynein-Lis1), and Ndel1 and Lis1 are always bound in these experiments given their high-affinity interaction (Figure 1C; Figure S4G; Figure 4A). Here, Ndel1 would be bound to Lis1 and dynein simultaneously without Lis1 engaging the motor at the AAA3/4 and stalk sites (Figure 4K).

To differentiate between the two possible binding modes, we employed FL-Ndel1^E48A^, which disrupts binding to dynein intermediate chain. If the dynein-Lis1-Ndel1 complex is tripartite, FL-Ndel1^E48A^ would reduce the Lis1-dynein binding to the levels observed when no Ndel1 was included (∼40%). However, if Lis1 cannot bind dynein and Ndel1 at the same time and the increased affinity is due to a Ndel1 daisy-chain, then FL-Ndel1^E48A^ would inhibit Lis1-dynein binding. When we performed the experiment with FL-Ndel1^E48A^, we observed almost no dynein depletion by Lis1 (Figure 4I). This result supports the daisy-chain model (Figure 4K) and suggests that Lis1 cannot bind Ndel1 and dynein at the same time. This finding is consistent with a recent study that also showed that Lis1 does not bind Ndel1 and dynein concurrently [49]. The daisy-chain model of Lis1-dynein-Ndel1 binding also explains the dominant effect of FL-Ndel1^PM^ over Lis1 in the landing rate of DDB measured in the smTIRF motility experiments. The Lis1 residues that contribute to an interaction with dynein’s stalk are very close to residues that mediate interaction with the Ndel1 orthologue, Nde1 (Figure S4H) [21, 67, 68], suggesting an overlapping binding site. Additionally, in a recent study, Okada and colleagues observe that mutations in Lis1 that disrupt binding to AAA3/4 abrogate Ndel1-Lis1 association, which also supports the hypothesis that Lis1 uses a close or overlapping site to bind Ndel1 and dynein [49].

## Discussion

We set out to explore how the core dynein regulator, Ndel1, affects activation of dynein motility. We found that Ndel1’s C-terminal half functions in a regulatory capacity to modulate binding to both dynein and Lis1. First, we confirmed that Ndel1’s C-terminal coiled-coil participates in dynein binding (Figure 1C) and found that phosphomimetic mutations in the C-terminus that mimic CDK5 phosphorylation positively regulate binding to dynein (Figure 3B). Because many kinases target similar residues as CDK5, we propose that phosphorylation by multiple kinases may promote increased dynein-Ndel1 interaction (Figure S3A). Further work is needed to understand how Ndel1’s C-terminal coiled-coil promotes dynein binding and to determine the structural effect of phosphorylation. Our finding that Ndel1’s C-terminal half negatively regulates association with Lis1 (Figure 4A) suggests that FL-Ndel1 is autoinhibited with respect to Lis1 binding. Structural predictions and previous studies suggest that Ndel1’s C-terminus contains a stretch of amino acids that can exist as a coiled-coil or fold back and interact with the interface that Ndel1 uses to bind Lis1 (Figure S1B, C; Figure 4B; Figure S4D) [27, 69]. We posit that Ndel1 may exploit different conformational states to regulate association with Lis1 (Figure S4E). More studies will be required to test this hypothesis.

Our work also shows that Ndel1 directly disfavors the formation of the activated dynein complex and that the N-terminal coiled-coil is necessary and sufficient to mediate this effect (Figure 1D-I). Previous reports have shown that Ndel1 and the p150 subunit of dynactin compete for binding to the intermediate chain N-terminus because they have overlapping, yet non-identical binding sites [46, 48]. Although the p150-intermediate chain interface is not resolved in any published structures of dynein-dynactin-adaptor, a recent study shows that the interaction between p150 and the intermediate chain promotes or stabilizes the activated dynein complex [49]. Therefore, we suggest that Ndel1 disfavors complex formation by competing with p150 for intermediate chain binding. We found that Ndel1’s C-terminal coiled-coil promotes binding to dynein and phosphomimetic mutations in the C-terminus can amplify the inhibitory effect (Figure 1C-F; Figure 3B-F). Therefore, we suggest that the C-terminal half of Ndel1 acts to stabilize the interaction with dynein and provide a mechanism for the magnitude of inhibition to be modulated by phosphorylation. Exploring the role of the p150-intermediate chain interaction in activated complex formation will be critical to understand how Ndel1 fits into the current model of dynein activity. We also observe that Ndel1 inhibits the formation of activated dynein complexes to different extents, depending on the identity of the adaptor (Figure 2). This is likely because different adaptors form activated complexes with varying stabilities. This result suggests that Ndel1-dynein interaction in cells may have a different outcome depending on the adaptor present.

Additionally, our work shows that Ndel1 also inhibits dynein activation by preventing the direct association of dynein and Lis1 (Figure 4I). We observe that though Ndel1 can bind dynein and Lis1 at the same time, Lis1 does not bind both Ndel1 and dynein concurrently. This competition is likely because dynein and Ndel1 have close or overlapping binding sites on Lis1. Structural studies will be required to fully test this theory.

Together, our data shows that Ndel1 operates in two distinct ways to disfavor dynein activation: directly by preventing association of the dynein-dynactin-adaptor complex and indirectly by inhibiting the activator Lis1. However, most published cell-based studies suggest that Ndel1 works with Lis1 to positively regulate dynein activation. How can we reconcile our findings with what has been observed in cells? Given that Ndel1 also promotes dynein localization to key subcellular locations, we propose that Ndel1 acts as a scaffold for dynein and Lis1 (Figure 5, step 1). In this role, we hypothesize that Ndel1 drives dynein-Lis1 co-localization, while simultaneously holding dynein in an inactive state by preventing dynactin binding and Lis1 activity (Figure 5, step 1-3). Because the FL-Ndel1^PM^ was a more potent inhibitor of activated dynein complex assembly, we propose that phosphorylation of Ndel1’s C-terminal domain also regulates its ability to recruit and inhibit dynein. In this model, Ndel1 must unbind from dynein and Lis1 for dynein activation to occur (Figure 5, step 4, 5). We speculate that dephosphorylation of Ndel1 may aid in dissociation from dynein given the increased affinity of the FL-Ndel1^PM^-dynein interaction, however it is possible that there are other mechanisms to cause Ndel1 release. For example, at the kinetochore Ndel1 may interact with CENP-F in a manner that attenuates dynein’s ability to traffic checkpoint proteins [29, 30]. It’s possible that CENP-F-Ndel1 dissociation may promote Ndel1-dynein dissociation, and thus dynein activation. We also propose that conformational changes in Ndel1 where the C-terminal tail folds back may facilitate Lis1 release (Figure 5, step 5). How this conformational change would be triggered is currently unknown. Once uncoupled from Ndel1, dynein activation can proceed (Figure 5, states 6, 7). It is possible that Ndel1 may also help “reset” dynein to an inactive state by destabilizing the activated dynein complex.

**Figure 5.**
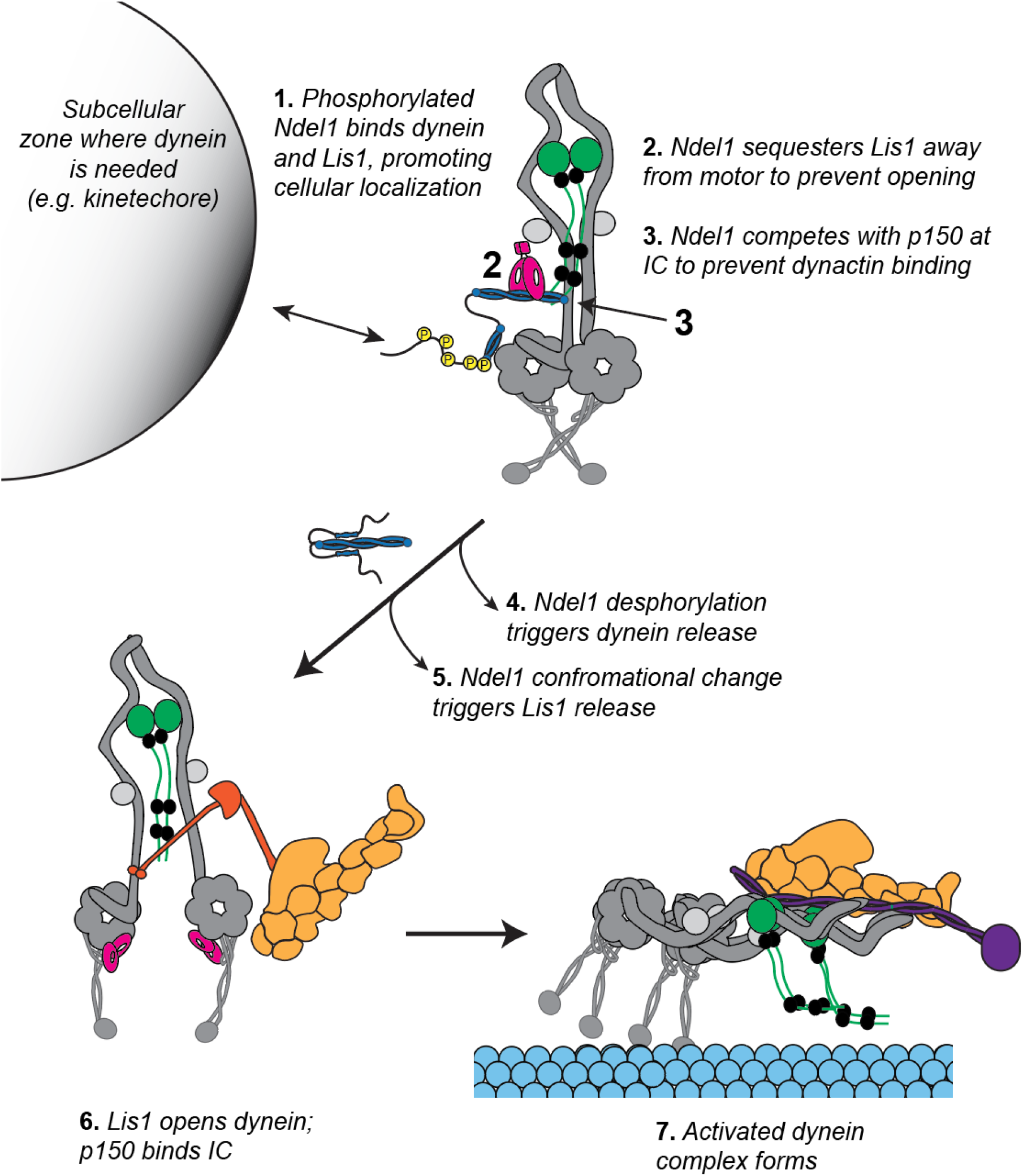
Model of how Ndel1 regulates the formation of activated dynein complexes through its interaction with dynein and Lis1. Step 1) Ndel1 and its orthologue Nde1 stabilize dynein and Lis1 at multiple cellular regions, including the kinetochore and nuclear pore [10, 29, 61]. Phosphorylation by CDK5 and/or CDK1 (which target nearly identical amino acids in Ndel1) are critical for Ndel1-mediated recruitment of dynein and Lis1 [61, 62]. **Steps 2 and 3)** Ndel1 simultaneously binds Lis1 and sequesters it from its AAA3/4 and stalk binding sites on dynein and inhibits the interaction between p150 and the intermediate chain, thus holding dynein in an inactive state. **Steps 4 and 5)** Ndel1 dissociation from Lis1 and dynein is required for dynein activation. This may be controlled by dephosphorylation and a conformational change, which reduces affinity for dynein and Lis1, respectively. How dephosphorylation or conformational change would be triggered is currently unknown. **Steps 6 and 7)** Dynein activation can proceed as has been described [14–16, 19–21, 25, 78]. Our data suggests that p150-intermediate chain interaction is critical for formation of the activated dynein complex.

Why would a scaffold that holds dynein in an inhibited state ultimately support dynein activation? It’s possible that Ndel1-bound dynein and Lis1 are “primed” for activation, forming a structure that is more amenable to activated dynein complex formation than without Ndel1. If this is true, then Ndel1 holds Lis1 and dynein in such a way to increase the effectiveness of Lis1-mediated activated dynein complex formation. It is also possible that Ndel1 promotes activation of dynein by simply increasing the colocalization and thus effective concentrations of dynein and Lis1.

It is intriguing that Ndel1 holds dynein in an inactive state and promotes its localization. Being able to recruit populations of inhibited dynein to cellular structures would enable a concerted activation of all dynein in a given cellular region. This would be particularly important for events where dynein activation must be precisely timed, like during mitosis. For example, to support the metaphase-anaphase transition, dynein traffics Spindle Assembly Checkpoint proteins away from the kinetochore. Here, both premature and delayed activation of dynein would be disastrous, leading to chromosome missegregation [73]. Additionally, dynein activation needs to be synchronized across all kinetochores of the cell to facilitate proper chromosome segregation [74]. We speculate that Ndel1 release of dynein can function as a timed trigger to promote dynein activation en masse. More work is required to test this model and establish the cellular signals that modulate Ndel1-dynein association.

## Acknowledgements

We thank members of the DeSantis lab, Ryoma Ohi, Michael Cianfrocco, Jayakrishnan Nandakumar, Kristen Verhey, David Sept, Richard McKenney, Steven Markus, Ahmet Yildiz, and Samara Reck-Peterson for helpful discussions and feedback. We thank the Lavis lab and Janelia Research Campus for sharing Halo-JFX646 dye. Thank you to Ricardo Henriques for the bioRxiv template. This work was supported by NIH grants R00GM127757 (M.D.) and R35GM146739 (M.D.) and NSF grant 2142670 (M.D.).

## Author Contributions

S.R.G, J.P.G., and M.E.D. conceived the study. S.R.G., J.P.G., and A.S. purified proteins for the study. S.R.G., J.P.G., A.S., S.R.L., and R.E. performed smTIRF experiments and analyzed data. S.R.G. and J.P.G. performed pulldown experiments. All authors contributed to the writing of the manuscript.

## Author Information

The authors declare no competing financial interests.

## Methods

### Constructs

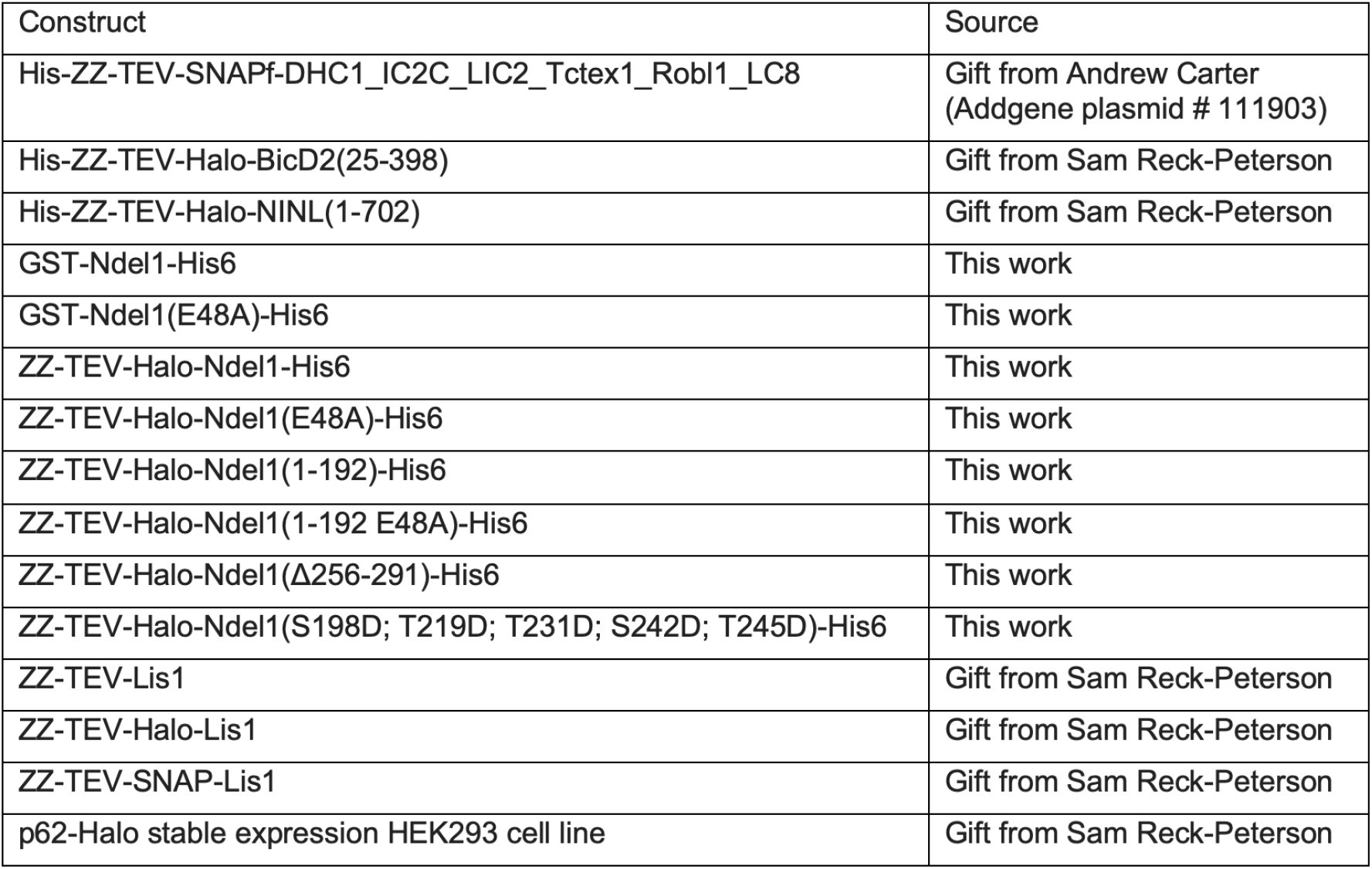

#### Protein Expression and Purification

Dynein, Lis1, and Ndel1 constructs were expressed in Sf9 cells as described [2, 75]. Briefly, pACEBac1 plasmid containing the human dynein genes, pFastBac plasmid containing full-length Lis1, and pKL plasmid containing Ndel1 and tagged Lis1 constructs were transformed into DH10EmBacY chemically competent cells with heat shock at 42°C for 15 s followed by incubation at 37°C and shaking at 220 rpm for 6 h in S.O.C. media (Thermo Fisher Scientific). The cells were plated on LB-agar plates containing kanamycin (50 µg/mL), gentamicin (7 µg/mL), tetracycline (10 µg/mL), BluoGal (100 µg/mL) and IPTG (40 µg/mL). Cells that contained the plasmid of interest were identified with blue/white selection after 48-72 h. For full-length human dynein constructs, white colonies were tested for the presence of all six dynein genes with PCR. Colonies were grown overnight in LB medium containing kanamycin (50 µg/mL), gentamicin (7 µg/mL) and tetracycline (10 µg/mL) at 37°C and agitation at 220 rpm. Bacmid DNA was extracted from overnight cultures using isopropanol precipitation as described[14]. 1 × 106 Sf9 cells in 2 mL of media in a 6-well dish were transfected with up to 2 µg of fresh bacmid DNA using FuGene HD transfection reagent (Promega) at a ratio of 3:1 (Fugene reagent:DNA) according to the manufacturer’s directions. Cells were incubated at 27°C for 3 d without agitation in a humid incubator. Next, the supernatant containing the virus (V0) was harvested by centrifugation (1000 x g, 5 min, 4°C). 1 mL of the V0 virus was used to transfect 50 mL of Sf9 cells at 1 × 106 cells/mL to generate the next passage of virus (V1). Cells were incubated at 27°C for 3 d with shaking at 105 rpm. Supernatant containing V1 virus was collected by centrifugation (1000 x g, 5 min, 4°C). All V1 was protected from light and stored at 4°C until further use. To express protein, 4 mL of V1 virus were used to transfect 400 mL of Sf9 cells at a density of 1 × 106 cells/mL. Cells were incubated at 27°C for 3 d with shaking at 105 rpm and collected by centrifugation (3500 x g, 10 min, 4°C). The pellet was washed with 10 mL of ice-cold PBS and collected again via centrifugation before being flash-frozen in liquid nitrogen and stored at -80°C until needed for protein purification.

All steps for protein purification were performed at 4°C unless indicated otherwise. For dynein preparation, Sf9 cell pellets were thawed on ice and resuspended in 40 mL of Dynein-lysis buffer (50 mM HEPES (pH 7.4), 100 mM sodium chloride, 1 mM DTT, 0.1 mM Mg-ATP, 0.5 mM Pefabloc, 10% (v/v) glycerol) supplemented with 1 cOmplete EDTA-free protease inhibitor cocktail tablet (Roche) per 50 mL. Cells were lysed with a Dounce homogenizer (10 strokes with a loose plunger followed by 15 strokes with a tight plunger). The lysate was clarified by centrifugation (183,960 x g, 88 min, 4°C) in a Type 70Ti rotor (Beckman). The supernatant was mixed with 2 mL of IgG Sepharose 6 Fast Flow beads (Cytiva) equilibrated in Dynein-lysis buffer and incubated for 3-4 h with rotation along the long axis of the tube. The beads were transferred to a glass gravity column, washed with at least 200 mL of Dynein-lysis buffer and 300 mL of TEV buffer (50 mM Tris–HCl (pH 8.0), 250 mM potassium acetate, 2 mM magnesium acetate, 1 mM EGTA, 1 mM DTT, 0.1 mM Mg-ATP, 10% (v/v) glycerol). For fluorescent labeling of SNAP tag, dynein-bound beads were mixed with 5 µM SNAP-Cell-TMR or SNAP-AlexaFluor-647 (New England Biolabs) for 10 min at RT. Unconjugated dye was removed with a 300 mL wash with TEV buffer at 4°C. The beads were resuspended in 15 mL of TEV buffer supplemented with 0.5 mM Pefabloc and 0.2 mg/mL TEV protease and incubated overnight with rotation along the long axis of the tube. Cleaved proteins in the supernatant were concentrated with a 100K MWCO concentrator (EMD Millipore) to 500 µl and purified via size exclusion chromatography on a TSKgel G4000SWXL column (TOSOH Bioscience) with GF150 buffer (25 mM HEPES (pH 7.4), 150 mM KCl, 1 mM MgCl_2_, 1 mM DTT) supplemented with 0.1 mM Mg-ATP as the mobile phase at 0.75 mL/min. Peak fractions were collected, buffer exchanged into a GF150 buffer supplemented with 0.1 mM Mg-ATP and 10% glycerol, and concentrated to 0.1-0.5 mg/mL using a 100K MWCO concentrator (EMD Millipore). Small volume aliquots were flash frozen in liquid nitrogen and stored at -80°C.

Lysis and clarification steps for the Ndel1 and Lis1 purifications were similar to the dynein purification except Lis1-lysis buffer (30 mM HEPES (pH 7.4), 50 mM potassium acetate, 2 mM magnesium acetate, 1 mM EGTA, 300 mM potassium chloride, 1 mM DTT, 0.5 mM Pefabloc, 10% (v/v) glycerol supplemented with 1 cOmplete EDTA-free protease inhibitor cocktail tablet (Roche) per 50 mL) was used in place of dynein-lysis buffer. The clarified supernatant was mixed with 2 mL of IgG Sepharose 6 Fast Flow beads (GE Healthcare Life Sciences) and incubated for 2-3 h with rotation along the long axis of the tube. The beads were transferred to a gravity column, washed with at least 20 mL of Lis1-lysis buffer, 200 mL of Lis1-TEV buffer (10 mM Tris–HCl (pH 8.0), 2 mM magnesium acetate, 150 mM potassium acetate, 1 mM EGTA, 1 mM DTT, 10% (v/v) glycerol) supplemented with 100 mM potassium acetate and 0.5 mM Pefabloc; and 100 mL of Lis1-TEV buffer. For fluorescent labeling of Ndel the Ndel-bound beads were mixed with 5 µM Halo-JFX646 (Lavis Lab) for 10 min at RT after the lysis buffer wash. TEV protease was added to the beads at a final concentration of 0.2 mg/mL, and the beads were incubated overnight with rotation along the long axis of the tube. Cleaved Lis1 or Ndel1 in the supernatant was collected and concentrated to 500 µl with a 30K MWCO concentrator (EMD Millipore). The concentrated protein was then purified via size exclusion chromatography on a Superose 6 Increase 10/300 GL column (Cytiva) with GF150 buffer as the mobile phase at 0.75 mL/min. For Lis1 the GF150 mobile phase was supplemented with 10% glycerol. SNAP-Lis1 was labelled with SNAP-AlexaFluor 647 (Promega) by incubating with 5uM dye for 10min at RT prior to size exclusion chromatography. Peak fractions were collected, concentrated to 0.2-1 mg/mL with a 30K MWCO concentrator (EMD Millipore), frozen in liquid nitrogen, and stored at -80°C. Ndel1 purifications were supplemented with 10% glycerol before freezing.

Dynactin was purified from HEK 293T cell lines stably expressing p62-Halo-3xFLAG as described[58]. Briefly, frozen pellets collected from 160 15 cm plates were resuspended in 80 mL of Dynactin-lysis buffer (30 mM HEPES (pH 7.4), 50 mM potassium acetate, 2 mM magnesium acetate, 1 mM EGTA, 1 mM DTT, 10% (v/v) glycerol) supplemented with 0.5 mM Mg-ATP, 0.2% Triton X-100, and 1x cOmplete EDTA-free protease inhibitor cocktail tablets (Roche)) and rotated along the long axis of the tube for at least 15 min. The lysate was clarified via centrifugation (66,000 x g, 30 min, 4°C) in a Type 70 Ti rotor (Beckman). The supernatant was mixed with 1.5 mL of anti-FLAG M2 affinity gel (Sigma-Aldrich) and incubated overnight with rotation along the long axis of the tube. The beads were transferred to a glass gravity column, washed with at least 50 mL of wash buffer (Dynactin-lysis buffer supplemented with 0.1 mM Mg-ATP, 0.5 mM Pefabloc and 0.02% Triton X-100), 100 mL of wash buffer supplemented with 250 mM potassium acetate, and then washed again with 100 mL of wash buffer. 1 mL of elution buffer (wash buffer with 2 mg/mL of 3xFlag peptide) was used to elute dynactin, which was then filtered via centrifuging through a Ultrafree-MC VV filter (EMD Millipore) in a tabletop centrifuge according to the manufacturer’s instructions. The filtered dynactin was then diluted to 2 mL in Buffer A (50 mM Tris-HCl (pH 8.0), 2 mM magnesium acetate, 1 mM EGTA, and 1 mM DTT) and loaded onto a MonoQ 5/50 GL column (Cytiva) at 1 mL/min. The column was pre-washed with 10 CV of Buffer A, 10 CV of Buffer B (50 mM Tris-HCl (pH 8.0), 2 mM magnesium acetate, 1 mM EGTA, 1 mM DTT, 1 M potassium acetate) and then equilibrated with 10 CV of Buffer A. A linear gradient was run over 26 CV from 35-100% Buffer B. Pure dynactin eluted between 75-80% Buffer B. Peak fractions were collected, pooled, buffer exchanged into a GF150 buffer supplemented with 10% glycerol, concentrated to 0.02-0.1 mg/mL using a 100K MWCO concentrator (EMD Millipore), aliquoted into small volumes, then flash-frozen in liquid nitrogen.

BicD2 and NINL containing amino-terminal HaloTags were expressed in BL-21[DE3] cells (New England Biolabs) at an optical density at 600 nm of 0.4–0.6 with 0.1mM IPTG for 16h at 18°C. Frozen cell pellets from a 1.5L culture were resuspended in 40mL of activating-adaptor-lysis buffer (30mM HEPES pH7.4, 50mM potassium acetate, 2mM magnesium acetate, 1mM EGTA, 1mM DTT and 0.5mM Pefabloc, 10% (v/v) glycerol) supplemented with 1x cOmplete EDTA-free protease inhibitor cocktail tablets (Roche) and 1mg/mL lysozyme. The resuspension was incubated on ice for 30min and lysed by sonication. The lysate was clarified by centrifuging at 66,000g for 30min in Type 70 Ti rotor (Beckman). The clarified supernatant was incubated with 2mL of IgG Sepharose 6 Fast Flow beads (Cytiva) for 2h on a roller. The beads were transferred into a gravity-flow column, washed with 100mL of activating-adaptor-lysis buffer supplemented with 150mM potassium acetate and 50mL of cleavage buffer (50mM Tris-HCl pH8.0, 150mM potassium acetate, 2mM magnesium acetate, 1mM EGTA, 1mM DTT, 0.5mM Pefabloc and 10% (v/v) glycerol). The beads were then resuspended and incubated in 15mL of cleavage buffer supplemented with 0.2mg/mL TEV protease overnight with rotation along the long axis of the tube. The supernatant containing cleaved proteins was concentrated using a 30kDa MWCO concentrator (EMD Millipore) to 1mL, filtered by centrifuging with Ultrafree-MC VV filter (EMD Millipore) in a tabletop centrifuge, diluted to 2mL in buffer A (30mM HEPES pH7.4, 50mM potassium acetate, 2mM magnesium acetate, 1mM EGTA, 10% (v/v) glycerol and 1mM DTT) and injected into a MonoQ 5/50 GL column (Cytiva) at 1m/min. The column was prewashed with 10 CV of buffer A, 10 CV of buffer B (30mM HEPES pH7.4, 1m potassium acetate, 2mM magnesium acetate, 1mM EGTA, 10% (v/v) glycerol and 1mM DTT) and again with 10 CV of buffer A. To elute, a linear gradient was run over 26 CV from 0–100% buffer B. The peak fractions containing Halo-tagged activating adaptors were collected and concentrated to using a 30kDa MWCO concentrator (EMD Millipore) to 0.2mL, diluted to 0.5mL in GF150 buffer and further purified using size-exclusion chromatography on a Superose 6 Increase 10/300GL column (Cytiva) with GF150 buffer at 0.5mL/min. The peak fractions were collected, buffer-exchanged into a GF150 buffer supplemented with 10% glycerol, concentrated to 0.2–1mg/mL using a 30kDa MWCO concentrator (EMD Millipore) and flash-frozen in liquid nitrogen.

#### Structure Prediction

To predict the structure of full length Ndel1 and the C-terminal half of Ndel1, we used the COSMIC2 web-platform to run CoLab Fold on full length Ndel1 or Ndel1 (amino acids 194-345) [70, 71].

#### Protein Binding Assays

The binding affinity of Ndel1 constructs for dynein and Lis1 was determined by coupling Ndel1 to 25 µL of Magne HaloTag Beads (Promega) in 2 mL Protein Lo Bind Tubes (Eppendorf) using the following protocol. Beads were washed twice with 1 mL of GF150 without ATP supplemented with 10% glycerol and 0.1% NP40. Ndel1 was diluted in this buffer to 0, 30, 60, 90, 120, 300 and 600 nM. 25 µL of diluted Ndel1 was added to the beads and gently shaken for one hour. 20 µL of supernatant were then analyzed via SDS-PAGE to confirm complete depletion of Ndel1. The Ndel1-conjugated beads were washed once with 1 mL GF150 with 10% glycerol and 0.1% NP40 and once with 1mL of binding buffer (30 mM HEPES [pH 7.4], 2 mM magnesium acetate, 1 mM EGTA, 10% glycerol, 1 mM DTT, 1 mg/mL casein, 0.1% NP40, 1mM ADP) supplemented with 36 mM KCl (or 150mM KCl for high salt Lis1 curves). 5 nM dynein or Lis1 was diluted in binding buffer, which resulted in binding buffer with 36 mM KCl (or 150mM KCl for high salt Lis1 curves). 25 µL of the dynein or Lis1 mixture was added to the beads pre-bound with Ndel1 and gently agitated for 45 minutes. After incubation 20 µL of the supernatant was removed, and 6.67 µL of NuPAGE® LDS Sample Buffer (4X) and 1.33 µL of Beta-mercaptoethanol was added to each. The samples were boiled for 5 minutes before running on a 4-12% NuPAGE Bis-Tris gel. Depletion was determined using densitometry in ImageJ. Binding curves were fit in Prism9 (Graphpad) with a nonlinear regression for one site binding with Bmax set to 1.

Experiments measuring binding between dynein, Lis1 and Ndel1 were performed with the same method and buffers as above. For the experiment with Lis1 on the beads 50nM of Halo-Lis1 was conjugated to the beads and incubated with 5nM dynein with and without 100nM of WT and E48A Ndel1. For the experiment with Ndel1 on the beads 30nM of Halo-Ndel1 was conjugated to beads and incubated with 5nM dynein in the presence and absence of 30nM Lis1.

For experiments investigating the competition between Ndel1 and dynactin/activating adaptor the methods and buffers were the same as above. 90nM Ndel1 was conjugated to 30µL of NEBExpress® Ni-NTA Magnetic Beads (NEB) and incubated with 5nM dynein. The appropriate samples were further supplemented with 10nM dynactin and 150nM of wither BicD2 or NINL.

#### Single-molecule TIRF microscopy

Single-molecule imaging was performed with an inverted microscope (Nikon, Ti2-E Eclipse) with a 100x 1.49 N.A. oil immersion objective (Nikon, Apo). The microscope was equipped with a LUNF-XL laser launch (Nikon), with 405 nm, 488 nm, 561 nm, and 640 nm laser lines. The excitation path was filtered using an appropriate quad bandpass filter cube (Chroma). The emission path was filtered through appropriate emission filters (Chroma) located in a high-speed filter wheel (Finger Lakes Instrumentation). Emitted signals were detected on an electron-multiplying CCD camera (Andor Technology, iXon Ultra 897). Image acquisition was controlled by NIS Elements Advanced Research software (Nikon).

Single-molecule motility and microtubule binding assays were performed in flow chambers assembled as described previously[75, 76]. Either biotin-PEG-functionalized coverslips (Microsurfaces) or No. 1-1/2 coverslips (Corning) sonicated in 100% ethanol for 10 min were used for the flow-chamber assembly. Taxol-stabilized microtubules with ∼10% biotin-tubulin and ∼10% Alexa488 labelled fluorescent-tubulin were prepared as described[37]. Flow chambers were assembled with taxol-stabilized microtubules by incubating sequentially with the following solutions, interspersed with two washes with assay buffer (30 mM HEPES [pH 7.4], 2 mM magnesium acetate, 1 mM EGTA, 10% glycerol, 1 mM DTT) supplemented with 20 µM Taxol in between: (1) 1 mg/mL biotin-BSA in assay buffer (3 min incubation); (2) 0.5 mg/mL streptavidin in assay buffer (3 min incubation) and (3) a fresh dilution of taxol-stabilized microtubules in assay buffer (3 min incubation). After flowing in microtubules, the flow chamber was washed twice with assay buffer supplemented with 1 mg/mL casein and 20 µM Taxol.

To assemble dynein-dynactin-activating adaptor complexes, purified dynein (10-20 nM concentration), dynactin and the activating adaptor were mixed at 1:2:10 molar ratio and incubated on ice for 10 min. These dynein-dynactin-activating adaptor complexes were then incubated with Ndel1 and/or Lis1 or modified TEV buffer (to buffer match for experiments without Ndel1 or Lis1) for 10 min on ice. Dynactin and the activating adaptors were omitted for the experiments with dynein alone. The mixtures of dynein, dynactin, activating adaptor or dynein alone and Nde1l/Lis1 were then flowed into the flow chamber assembled with taxol-stabilized microtubules. The final imaging buffer contained the assay buffer supplemented with 20 µM Taxol, 1 mg/mL casein, 71.5 mM β-mercaptoethanol, 0.05 mg/mL glucose catalase, 1.2 mg/mL glucose oxidase, 0.4% glucose, and 2.5 mM Mg-ATP. The final concentration of dynein in the flow chamber was 0.5-1 nM for experiments with dynein-dynactin-activating adaptor complexes and 0.05 nM for dynein alone experiments. For standard motility experiments our final imaging buffer contained 37.5 mM KCl.

For single-molecule motility assays, microtubules were imaged first by taking a single-frame snapshot. Dynein labeled with fluorophores (TMR or Alexa647) was imaged every 300 msec for 3 min. At the end, microtubules were imaged again by taking a snapshot to assess stage drift. Movies showing significant drift were not analyzed. Each sample was imaged no longer than 10 min. For single-molecule microtubule binding assays, the final imaging mixture containing dynein was incubated for an additional 5 min in the flow chamber at room temperature before imaging. After 5 min incubation, microtubules were imaged first by taking a single-frame snapshot. Dynein labeled with fluorophores (TMR or Alexa64) were imaged by taking a single-frame snapshot. Each sample was imaged at 2 different fields of view and there were between 15 and 25 microtubules in each field of view. In order to compare the effect of Lis1 on microtubule binding, the samples with and without Lis1 were imaged in two separate flow chambers made on the same coverslip on the same day with the same stock of polymerized tubulin.

Kymographs were generated from motility movies, and dynein velocity and run length were calculated from kymographs using custom ImageJ macros as described [77]. Only runs longer than 8 pixels were included in the analysis. Bright protein aggregates, which were defined as molecules 4x brighter than the background, were excluded. Velocity and run length were reported for individual dynein complexes. The processive landing rate was reported per microtubule by dividing the total number of processive events analyzed for that microtubule by the length of the microtubule, divided by the concentration of dynein used in that experimental condition, divided by the length of the movie in minutes. Data plotting and statistical analyses were performed in Prism9 (GraphPad).

## Supplementary Materials

**Figure S1. Supplement to Figure 1.**
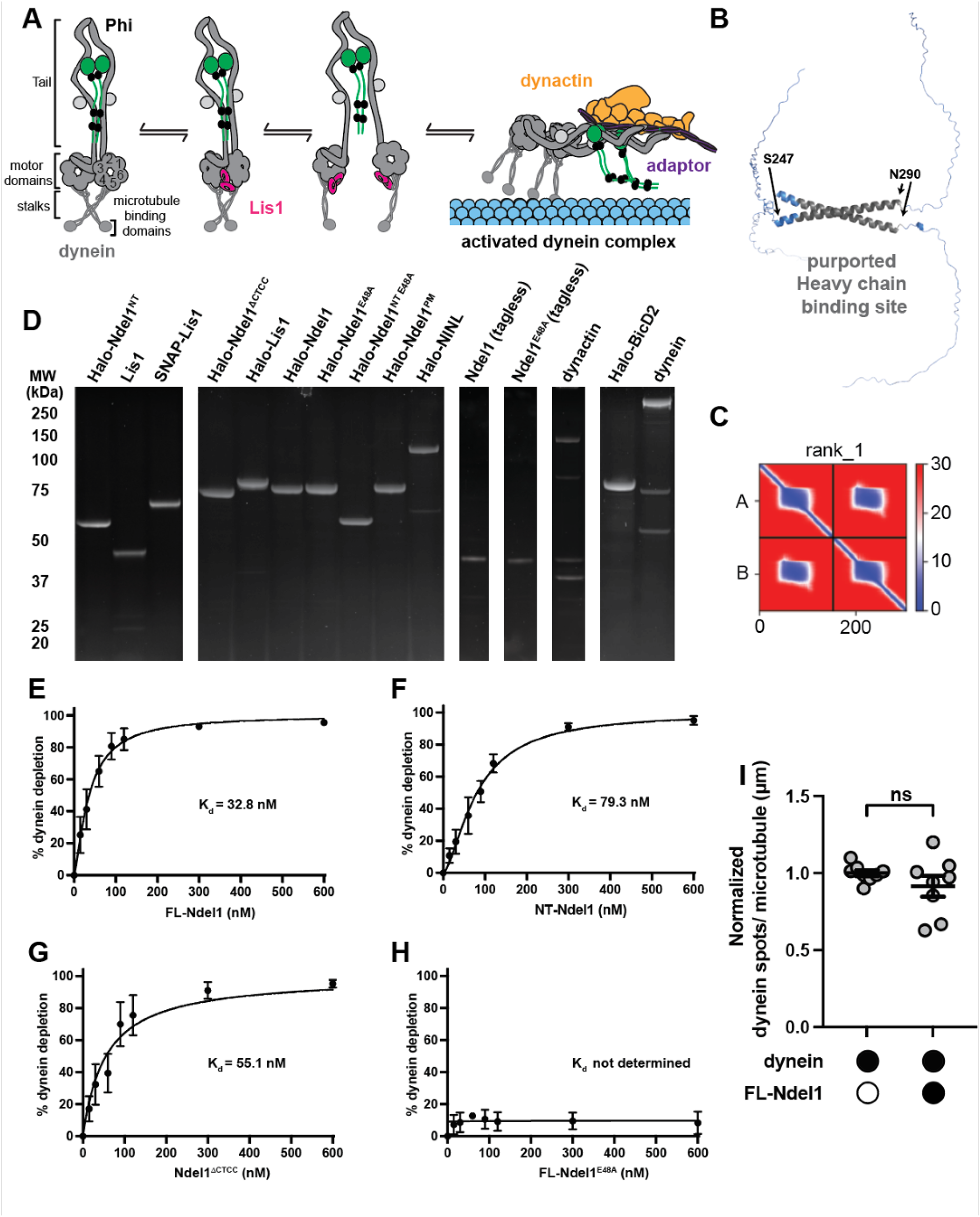
**A**. Model of Lis1 favoring open dynein and leading to activated dynein complex formation. **B**. Model of the C terminus of Ndel1 (from amino acids 194-345) generated in ColabFold showing alpha helical structure. Amino acids located at the beginning and end of the coiled-coil are indicated. Amino acids that have been shown to contribute to heavy chain binding are indicated in grey. **C**. Predicted alignment error plot for the ColabFold model in B. **D**. SDS-PAGE gels of all purified proteins used in this study. **E**. Binding curve between dynein and FL-Ndel1. n = 6. Error bars are mean ± SD. **F**. Binding curve between dynein and NT-Ndel1. n = 4. Error bars are mean ± SD. **G**. Binding curve between dynein and FL-Ndel1^ΔCTCC^. n = 3. Error bars are mean ± SD. **H**. Binding curve between dynein and FL-Ndel1^E48A^. n = 3. Error bars are mean ± SD. **I**. Normalized dynein spots/µm in the absence (white circles) and presence (black circles) of 300nM FL-Ndel1. Each point represents a field of view with 15-25 microtubules. n = 8. Statistical analysis was performed using Welch’s T test. P values: ns = 0.2641.

**Supplement to Figures 1 and 2.**
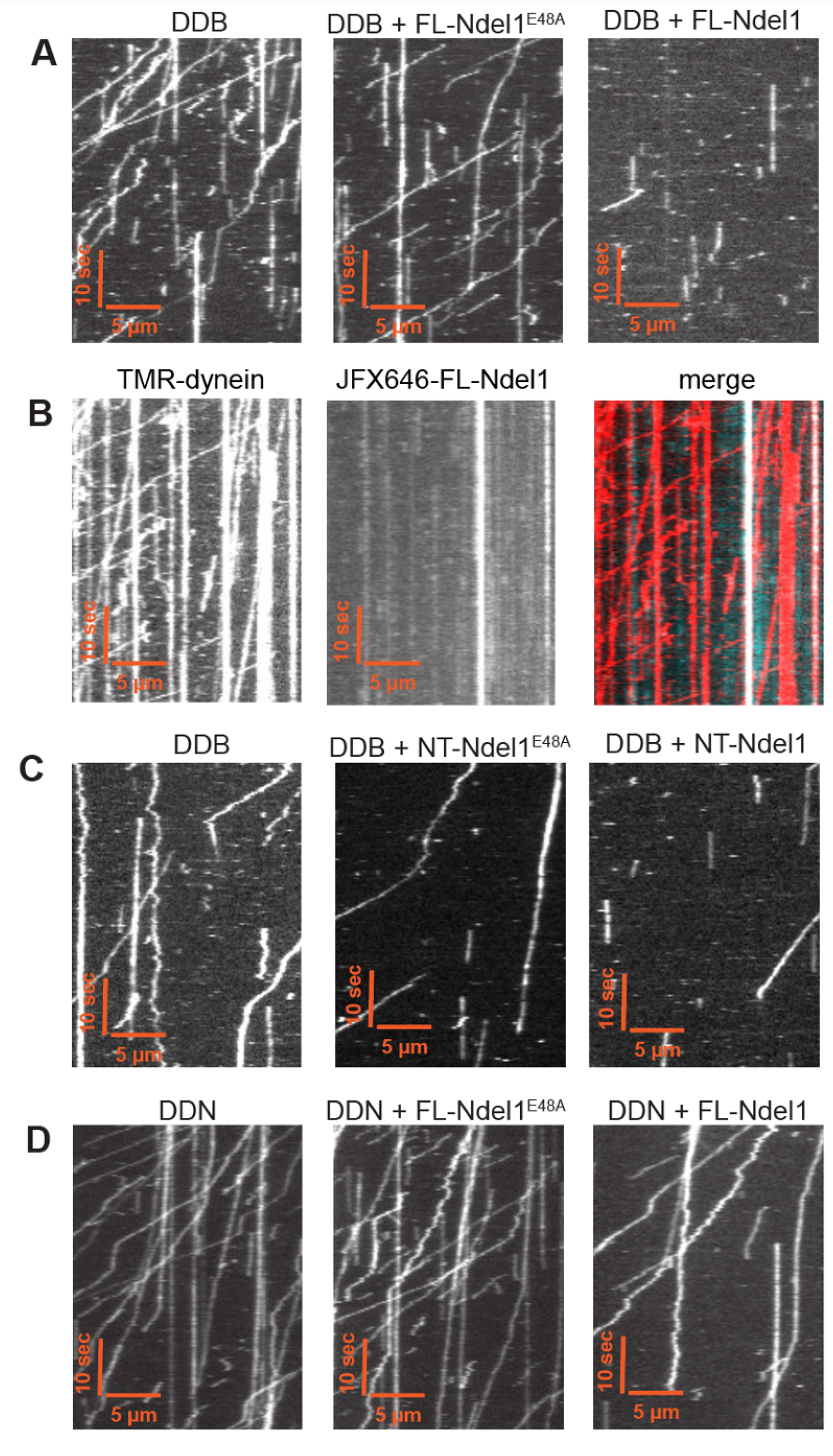
**A**. Example kymographs of DDB with and without 300nM FL-Ndel1^E48A^ and FL-Ndel1. **B**. Example kymographs of DDB (TMR dynein; red in merge) with 10nM Janelia Fluor-647 labelled FL-Ndel1 (cyan in merge). **C**. Example kymographs of DDB with and without 300nM NT-Ndel1^E48A^ and NT-Ndel1. **D**. Example kymographs of DDN with and without 300nM FL-Ndel1^E48A^ and FL-Ndel1.

**Figure S3. Supplement to Figure 3.**
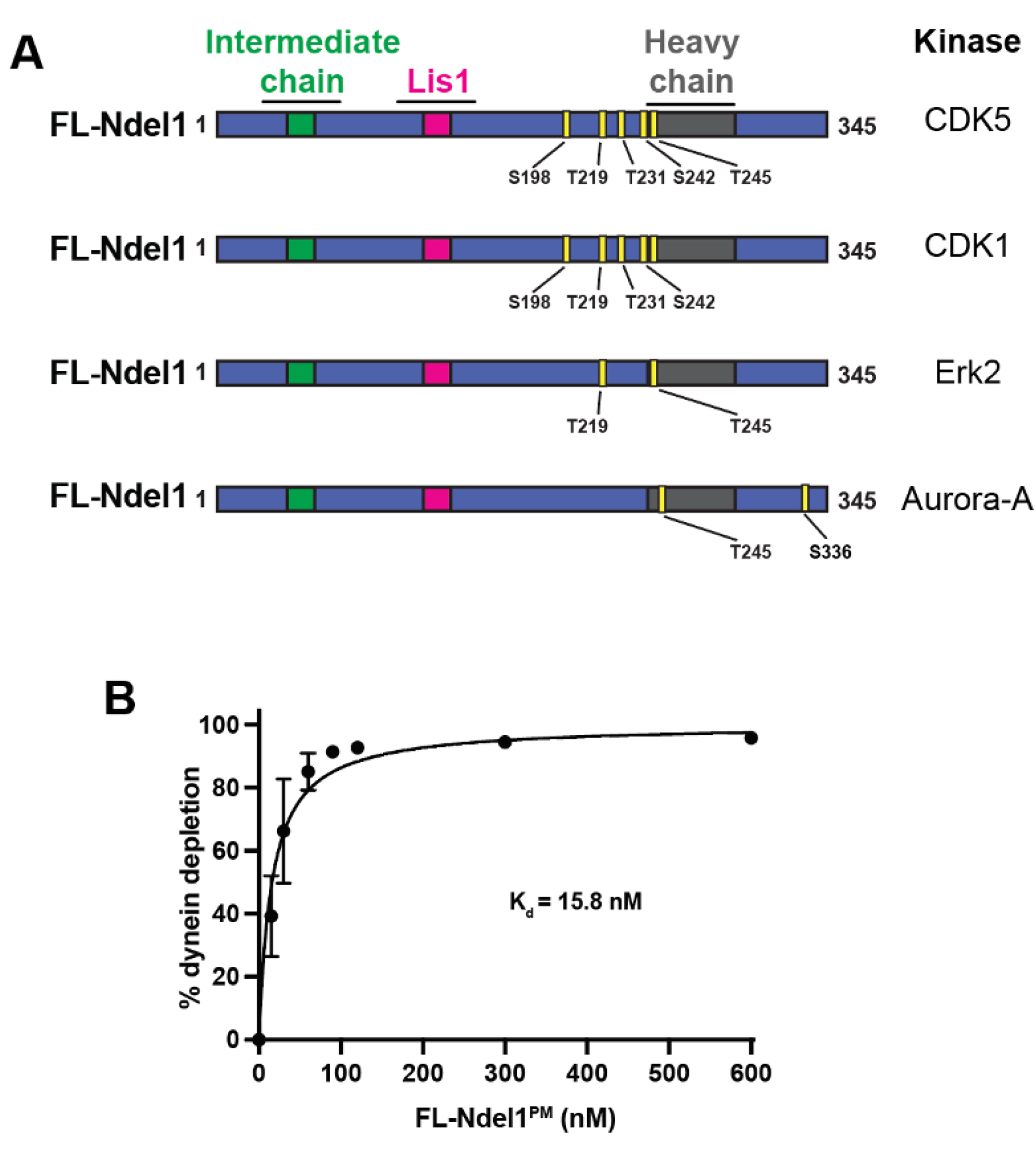
**A**. Schematics of FL-Ndel1 showing the residues phosphorylated by CDK5, CDK1, Erk2 and Aurora-A. **B**. Binding curve between dynein and FL-Ndel1^PM^. n = 3. Error bars are mean ± SD.

**Figure S4. Supplement to Figure 4.**
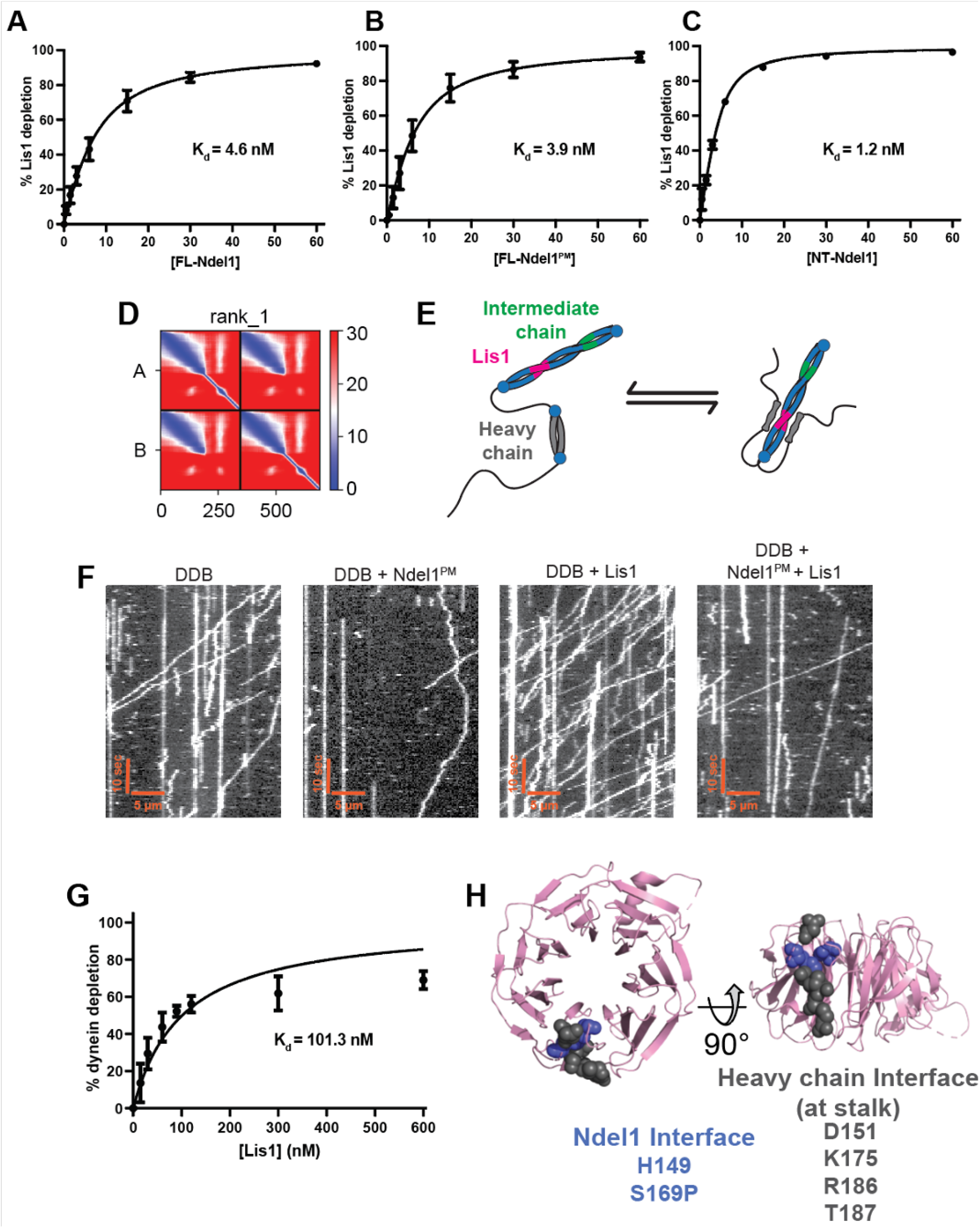
**A**. Binding curve between Lis1 and FL-Ndel1. n = 6. Error bars are mean ± SD. **B**. Binding curve between Lis1 and FL-Ndel1^PM^. n = 4. Error bars are mean ± SD. **C**. Binding curve between Lis1 and NT-Ndel1. n = 3. Error bars are mean ± SD. **D**. Predicted alignment error plot for the ColabFold model in Figure 4B. **E**. Conformational equilibrium of FL-Ndel1. **F**. Example kymographs of DDB with and without 50nM FL-Ndel1^PM^ and Lis1. **G**. Binding curve between dynein and Lis1. N = 3. Error bars are mean ± SD. **H**. H. Model of the Lis1 beta propeller (pink) (PDB: 1VYH) showing residues known to interact with Ndel1(blue) and dynein’s stalk (gray) [21, 67, 68].

